# A bacterial TIR-based immune system senses viral capsids to initiate defense

**DOI:** 10.1101/2024.07.29.605636

**Authors:** Cameron G. Roberts, Chloe B. Fishman, Dalton V. Banh, Luciano A. Marraffini

## Abstract

Toll/interleukin-1 receptor (TIR) domains are present in immune systems that protect prokaryotes from viral (phage) attack. In response to infection, TIRs can produce a cyclic adenosine diphosphate-ribose (ADPR) signaling molecule, which activates an effector that depletes the host of the essential metabolite NAD+ to limit phage propagation. How bacterial TIRs recognize phage infection is not known. Here we describe the sensing mechanism for the staphylococcal Thoeris defense system, which consists of two TIR domain sensors, ThsB1 and ThsB2, and the effector ThsA. We show that the major capsid protein of phage Φ80α forms a complex with ThsB1 and ThsB2, which is sufficient for the synthesis of 1’’-3’ glycocyclic ADPR (gcADPR) and subsequent activation of NAD+ cleavage by ThsA. Consistent with this, phages that escape Thoeris immunity harbor mutations in the capsid that prevent complex formation. We show that capsid proteins from staphylococcal Siphoviridae belonging to the capsid serogroup B, but not A, are recognized by ThsB1/B2, a result that suggests that capsid recognition by Sau-Thoeris and other anti-phage defense systems may be an important evolutionary force behind the structural diversity of prokaryotic viruses. More broadly, since mammalian toll-like receptors harboring TIR domains can also recognize viral structural components to produce an inflammatory response against infection, our findings reveal a conserved mechanism for the activation of innate antiviral defense pathways.

## INTRODUCTION

Numerous bacterial innate immune systems exhibit structural and/or functional homology with those observed in plants and animals (Wein and Sorek, 2022). In metazoans, the immune response begins with the recognition of pathogen-associated molecular patterns (PAMPs) by pattern recognition receptors such as the membrane-embedded Toll-like receptors (TLRs), leading in some cases to the synthesis of cyclic nucleotide second messengers that activate downstream effector responses (Burroughs et al., 2015; Doron et al., 2018; Li and Wu, 2021; Medzhitov et al., 1997). The Thoeris immune system of prokaryotes shares fundamental characteristics with TLR immune pathways, featuring proteins with toll/interleukin-1 receptor (TIR) domains capable of generating cyclic nucleotides that activate effector proteins to prevent viral (phage) propagation (Doron *et al*., 2018; Ka et al., 2020; Ledvina and Whiteley, 2024; Tamulaitiene et al., 2024). Present across nine distinct taxonomic phyla and found in approximately 4% of the bacterial and archaeal genomes examined, Thoeris systems are significantly represented in microbial defense mechanisms and can be classified into two different types depending on their genetic composition (Ofir et al., 2021). Type I Thoeris systems are comprised of a phage infection sensor with a TIR domain (ThsB) and an effector with a STALD (Sir2/TIR-Associating LOG-Smf/DprA) NAD-binding domain (ThsA). Upon viral infection, ThsB initiates the production of an nicotinamide adenine dinucleotide (NAD)-derived cyclic nucleotide, glycocyclic ADP-ribose (gcADPR) (Leavitt et al., 2022; Manik et al., 2022; Ofir *et al*., 2021), which binds to the STALD domain of ThsA and induces a conformational change that activates NAD+ degradation by the Sir2 domain (Ofir *et al*., 2021; Tamulaitiene *et al*., 2024). NAD+ depletion arrests the growth of the infected cells and prevents viral propagation, providing community-level immunity through the replication of the uninfected cells within the population (Ka *et al*., 2020).

How ThsB senses phage invasion to activate the type I Thoeris response remains unknown. Here we investigate this unanswered question in the bacterium *Staphylococcus aureus*. We demonstrate that the major head protein (Mhp), the most abundant component of viral capsids, from different staphylococcal phages directly interacts with ThsB TIR proteins to stimulate the synthesis of gcADPR and initiate the Thoeris immune response.

## RESULTS

### Thoeris provides anti-phage protection in staphylococci

A search for Thoeris systems in *S. aureus* revealed 32 unique but highly conserved operons (Supplementary Data File 1). We decided to investigate how the type I Thoeris system present in *S. aureus* 08BA02176 (Golding et al., 2012), hereafter designated Sau-Thoeris, senses phage infection to activate immunity. The Sau-Thoeris operon carries three genes (Fig. 1A) encoding two TIR-containing proteins, called ThsB1 and ThsB2, which produce a gcADPR signaling molecule (Leavitt *et al*., 2022; Ledvina and Whiteley, 2024; Ofir *et al*., 2021; Tamulaitiene *et al*., 2024), and a gcADPR-dependent effector, called ThsA, which was recently demonstrated to limit phage propagation by degrading NAD+ (Ofir *et al*., 2021; Tamulaitiene *et al*., 2024). To study this system, we cloned the full Sau-Thoeris operon (*thsA/B1/B2*), as well as different combinations of *ths* genes that we used as controls, into the staphylococcal vector pE194 (Horinouchi and Weisblum, 1982b) under the control of the P*spac* IPTG-inducible promoter for expression in the laboratory strain *Staphylococcus aureus* RN4220 (Kaltwasser et al., 2002). We tested immunity against seven different staphylococcal phages and found that expression of *thsA/B1/B2*, but not of *thsB1* or *thsB2* alone, reduced viral propagation, measured as plaque formation on top agar plates, of Φ80α-vir (Banh et al., 2023), ΦNM1γ6 (Goldberg et al., 2014), ΦNM4γ4 (Heler et al., 2015), ΦJ1, ΦJ2, and ΦJ4 (Banh *et al*., 2023), but not Φ12γ3 (Modell et al., 2017) (Fig. 1B). Similar results were obtained after infection of an *S. aureus* RN4220 strain carrying the Sau-Thoeris operon under the control of its native promoter in the chromosome (Fig. S1A).

**Figure 1.**
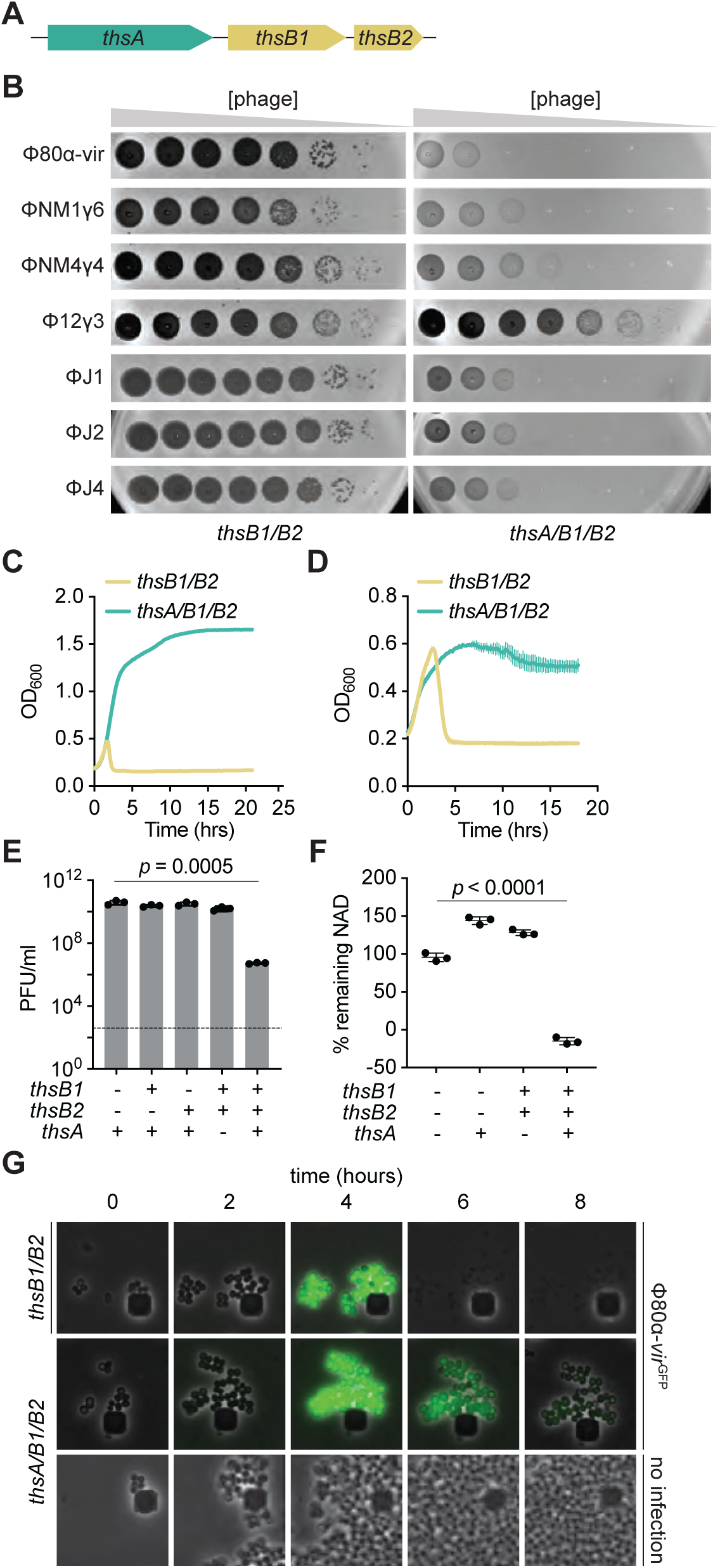
Thoeris provides anti-phage protection in staphylococci. **(A)** Schematic of the Thoeris operon present in the *Staphylococcus aureus* strain 08BA02176. The operon includes a *thsA* gene harboring a STALD domain, and two *thsB* genes, *thsB1* and *thsB2* that encode TIR domains. **(B)** Tenfold serial dilutions of different staphylococcal phages on lawns of *S. aureus* RN4220 harboring plasmids carrying either an incomplete (*thsB1/B2*) or full (*thsA/B1/B2*) Thoeris operon. **(C)** Growth of *S. aureus* RN4220 harboring plasmids carrying either an incomplete (*thsB1/B2*) or full (*thsA/B1/B2*) Thoeris operon, determined as the OD_600_ of the cultures after infection with Φ80α-vir at MOI 1. Mean of +/- S.D. of three biological replicates is reported. **(D)** Same as **(C)** but following the growth of lysogenic cultures after induction of the Φ80α prophage with MMC. **(E)** Enumeration of PFU/ml after induction of the Φ80α prophage with MMC present in lysogens harboring plasmids carrying different combinations of the *thsA*, *thsB1* and *thsB2* genes. Dotted line indicates the limit of detection. Mean of +/- S.D. of three biological replicates is reported; *p* value was obtained using an unpaired, two-tailed, *t*-test**. (F)** Measure of % remaining NAD+ and NADH (NAD), calculated as the ratio of the concentration of NAD+ and NADH detected in staphylococci harboring plasmids carrying different combinations of the *thsA*, *thsB1* and *thsB2* genes, to the value detected in the absence of any of the *ths* genes, after induction of the Φ80α prophage with MMC. Mean of +/- S.D. of three biological replicates is reported; *p* value was obtained using an unpaired, two-tailed, *t*-test**. (G)** Fluorescence microscopy of *S. aureus* RN4220 harboring plasmids carrying either an incomplete (*thsB1/B2*) or full (*thsA/B1/B2*) Thoeris operon. Images were taken every two hours after infection with Φ80α-vir-GFP phage, up to eight hours. The images are representative of three independent experiments.

Consistent with previous reports (Ofir *et al*., 2021), Sau-Thoeris immunity depended on the multiplicity of infection (MOI) and provided full defense of a bacterial culture at low phage concentrations (MOI 0.1 and 1; Figs. 1C and S1B), measured as the optical density at 600 nm (OD_600_) of the culture after infection. In addition, Sau-Thoeris immunity can be activated after the induction of a Φ80α prophage with the DNA-damaging agent mitomycin C (MMC), resulting in the growth of the bacterial culture (Fig. 1D) and in a significant decrease in the production of viable viral particles, measured as plaque-forming units (PFU) (Fig. 1E). We also measured activation of the Sau-Thoeris response using a colorimetric assay to quantify the depletion of oxidized and reduced NAD (NAD+ and NADH, respectively) in cell lysates upon induction of the Φ80α prophage, an experiment that resulted in a significant decrease in NAD detection after MMC treatment, only in the presence of the full *ths* operon (Fig. 1F). Finally, we used fluorescence microscopy to visualize Sau-Thoeris defense at the cellular level. We incubated staphylococci with a modified Φ80α-vir phage that expresses GFP (Φ80α-vir^GFP^) (Banh *et al*., 2023) in order to identify the infected cells. Green fluorescence decreased in staphylococci expressing ThsA/B1/B2, without cell lysis. In contrast, in cells harboring only the sensor genes *thsB1/B2*, GFP accumulation was followed by bacterial lysis (Fig. 1G). Altogether, these data show that, as previously reported for other species (Doron *et al*., 2018; Ofir *et al*., 2021), Sau-Thoeris activation results in a depletion of NAD+ levels that inhibits viral propagation.

### The phage major head protein activates Thoeris *in vivo*

After establishing a system to study Thoeris immunity in staphylococci, we set out to investigate how this response is triggered during phage infection. To do this we isolated Φ80α-vir phages that could form plaques in the presence of Sau-Thoeris, with the expectation that they would carry mutations in genes that are required for the activation of immunity. Since we were unable to observe discrete plaques after a single round of Φ80α-vir infection (Fig. 1B), we performed five sequential infections through the inoculation of supernatants of infected cultures into fresh staphylococci carrying Sau-Thoeris. We obtained individual plaques after this process, and we selected four to isolate the escaper phages and sequence their genomes (Supplementary Sequences 1). In all cases we detected the same mutation in *gp47*, the gene encoding Φ80α-vir’s major head protein (hereafter abbreviated Mhp for all the phages used in this study), which generated a missense amino acid substitution, V273A. To corroborate the role of this mutation in the evasion of Sau-Thoeris response, we infected liquid cultures with Φ80α-vir(*mhp*^V273A^). Addition of the mutant phage resulted in complete lysis in the presence of ThsA/B1/B2, and immunity was restored when staphylococci carried a second plasmid that expressed wild-type Mhp (Fig. 2A). Preparations of the escaper phage displayed the same plaquing efficiency and plaque size as wild-type Φ80α-vir in the absence of immunity (Fig. S2A). In the presence of Sau-Thoeris Φ80α-vir(*mhp*^V273A^) was able to form plaques, but at a lower efficiency than in the absence of defense. The viability of the escaper phage indicates that the V273A substitution does not prevent capsid formation and therefore it is possible that the mutation eludes immunity as part of a fully formed viral procapsid. To determine whether other proteins that are involved in capsid formation are required for the development of a proper Sau-Thoeris response, and to expand our results to a different staphylococcal virus, we performed deletions of the genes involved in capsid formation in the temperate phage ΦNM1 (Fig. S2B), through genetic engineering of prophages integrated in the genome of *S. aureus* RN4220. Capsid assembly in ΦNM1 begins with the formation of an empty precursor called the procapsid, comprised of 415 units of the major head protein (encoded by *gp43*) that directly associate with 100-200 units of a scaffolding protein (*gp42*), a 12-unit portal protein complex (*gp39*), which together with the terminase subunits is responsible for packaging the phage DNA into the capsid in an ATP-dependent manner (Quiles-Puchalt et al., 2014), and approximately 20 units of a minor head protein (*gp40*), whose role in capsid biogenesis is not clear (Spilman et al., 2011). This region (Fig. S2C) also harbors a small open reading frame of unknown function present, *gp41*. We knocked out these genes as well as *rinA*, required for the transcription of the phage structural genes (Ferrer et al., 2011), and complemented each deletion strain with plasmids expressing the missing gene, cloned on the staphylococcal plasmid pC194 (Horinouchi and Weisblum, 1982a) under the control of the IPTG-inducible P*spac* promoter (Kaltwasser *et al*., 2002). We induced lysogenic cultures with MMC and spotted the supernatants on lawns of staphylococci carrying an empty vector or a rescue plasmid (Fig. S2C). Except for *gp41*, we detected plaques only in the presence of the complementing vector, a result that corroborated both the essentiality of the disrupted genes as well as the identity of each mutant prophage. To test Sau-Thoeris activation, we induced the mutant prophages in the presence of *thsB1/B2* or *thsA/B1/B2* and measured NAD depletion (Fig. 2B). We found that, compared to a wild-type prophage control, induction of 11*rinA* and 11*gp43* mutants failed to reduce the levels of NAD within lysogens. Complementation of the Δ*gp43* lysogen with iis corresponding complementing plasmid (pMhp) restored NAD+ depletion after MMC treatment (Fig. 2C). Since absence of the portal, minor head or scaffolding proteins prevents the formation of the procapsid structure, these results demonstrate that the major head protein itself, and not a fully mature capsid, is necessary for the activation of the Sau-Thoeris response.

**Figure 2.**
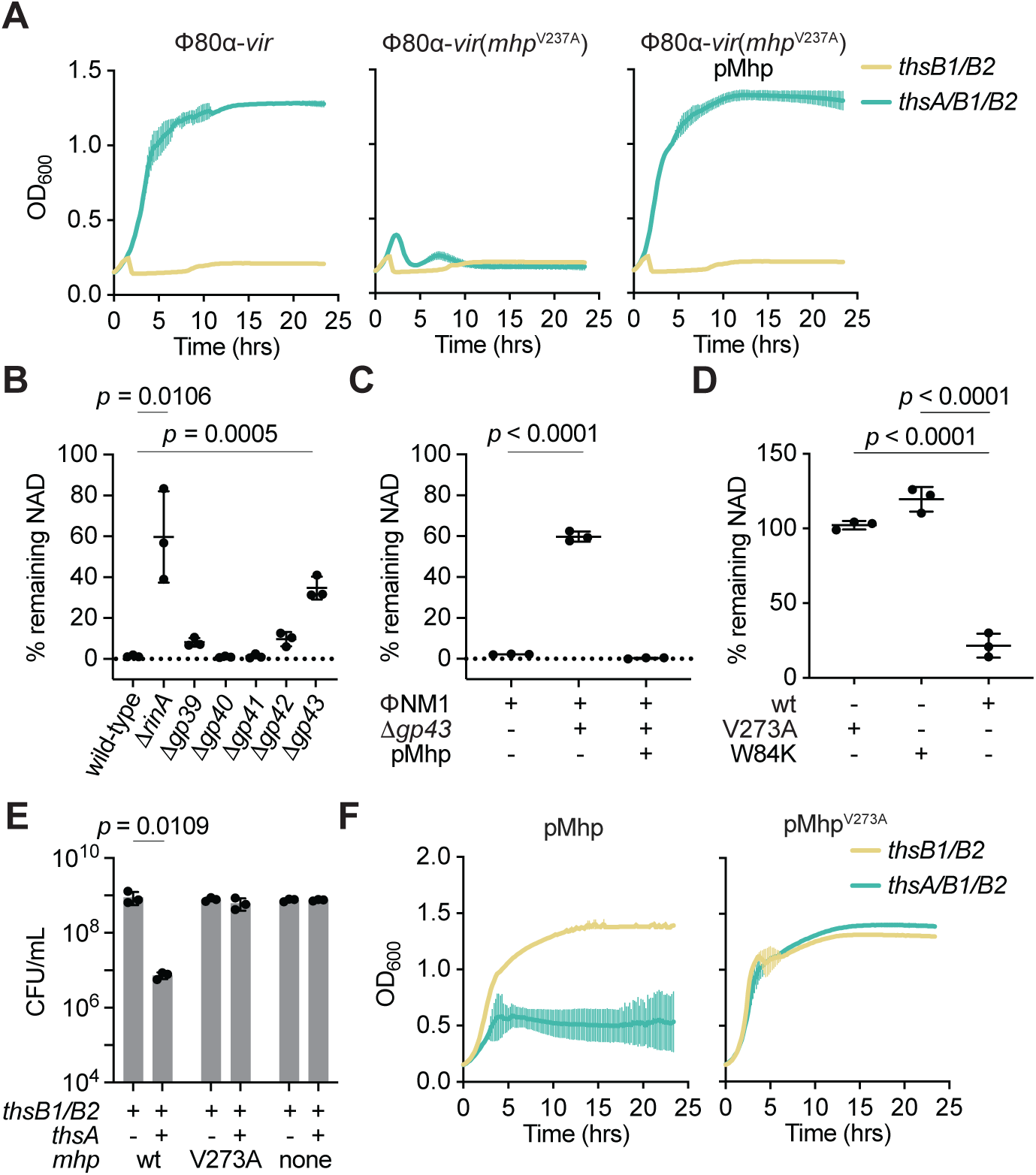
The phage major head protein activates Thoeris *in vivo*. **(A)** Growth of *S. aureus* RN4220 harboring plasmids carrying either an incomplete (*thsB1/B2*) or full (*thsA/B1/B2*) Thoeris operon in the absence or presence of a second plasmid expressing Mhp, determined as the OD_600_ of the cultures after infection with Φ80α-vir or Φ80α-vir(*mhp*^V273A^) at MOI 1. Mean of +/- S.D. of three biological replicates is reported. **(B)** Measure of % remaining NAD+/NADH (NAD), calculated as the ratio of the concentration of NAD+ and NADH detected in staphylococci harboring a plasmid carrying a full (*thsA/B1/B2*) Thoeris operon, to the value detected in the presence of an incomplete (*thsB1/B2*) system, after induction of the ΦNM1 prophage with MMC. Lysogens induced carried either wild-type or mutant prophages with deletions in different genes involved in capsid formation. Mean of +/- S.D. of three biological replicates is reported; *p* value was obtained using an unpaired, two-tailed, *t*-test**. (C)** Same as **(B)** after induction of wild-type and τι*gp43* ΦNM1 prophages, in the presence of a plasmid that expresses Mhp. **(D)** Same as **(B)** but after IPTG induction of expression of plasmid-encoded wild-type, V273A or W84K Mhp. **(E)** Enumeration of CFU/ml after induction of the Φ80α prophage with MMC present in lysogens harboring plasmids carrying either an incomplete (*thsB1/B2*) or full (*thsA/B1/B2*) Thoeris operon. Mean of +/- S.D. of three biological replicates is reported; *p* value was obtained using an unpaired, two-tailed, *t*-test**. (F)** Growth of *S. aureus* RN4220 harboring plasmids carrying either an incomplete (*thsB1/B2*) or full (*thsA/B1/B2*) Thoeris operon in the presence of a second plasmid expressing either wild-type or V273A Mhp, determined as the OD_600_ of the cultures after addition of IPTG. Mean of +/- S.D. of three biological replicates is reported.

To test if Mhp is also sufficient to trigger Sau-Thoeris immunity, we introduced pMhp into non-lysogenic *S. aureus* RN4220 carrying a second plasmid harboring different versions of the *ths* operon to achieve expression of the major head protein of ΦNM1 in the absence of phage infection. We measured NAD levels and found that only in the presence of the full *ths* operon, but not *thsB1/B2* alone, expression of Mhp significantly reduced the concentration of NAD within staphylococci (Fig. 2D). In contrast, introduction of the V273A escaper mutation into ΦNM1’s Mhp abrogated NAD depletion (Fig. 2D). Since NAD deficiency should inhibit the growth of staphylococci, which we observed during the microscopy analysis of infected hosts (Fig. 1G), we hypothesized that expression of wild-type, but not the V273A mutant, Mhp in the absence of phage infection should prevent the replication of staphylococci. Indeed, enumeration of colony-forming units (CFU) after addition of IPTG showed a significant decrease of viable cells in the presence of the full Thoeris operon when compared to the induction of cultures expressing only ThsB1/B2. In contrast IPTG induction of Mhp^V273A^ expression did not affect colony formation, showing a similar CFU count to that of cultures not expressing Mhp (Fig. 2E). In addition, growth of the cultures was also inhibited by over-expression of wild-type, but not V273A, Mhp (Fig. 2F). Altogether these results demonstrate that the major head protein of staphylococcal phages ΦNM1 and Φ80α is necessary and sufficient for the activation of the Sau-Thoeris response.

#### ThsB1/B2 form a complex with the major head protein *in vivo*

To determine whether the Mhp interacts directly with the Thoeris sensors, ThsB1 and ThsB2, during the phage lytic cycle, we constructed a plasmid that expressed both a hexahistidyl-tagged (His_6_) ThsB1 and 3xFLAG-tagged (FLAG) ThsB2, or the reverse, under the control of an IPTG-inducible promoter and introduced these constructs into *S. aureus* RN4220::ΦNM1 lysogens. We first confirmed that the addition of the tags to ThsB1 or ThsB2 did not impact function *in vivo* (Extended Data Fig. 3A). Following overexpression of tagged ThsB1 and ThsB2, we induced the prophage with MMC, and used a cobalt resin to separate the hexahistidyl-tagged ThsB1 or ThsB2. Western blot of the pulled-down proteins using anti-FLAG antibody showed the formation of a stable complex between ThsB1 and ThsB2 only when ΦNM1’s lytic cycle was induced, which depended on the presence of *gp43* (Fig. 3A). Importantly, SDS-PAGE of the pulled-down samples revealed the presence of a third protein that copurified with the ThsB1/B2 complex (Fig. 3B). Mass spectrometry analysis of the gel area containing this protein identified it as Mhp (Supplementary Data File 2 and Fig. S3B). We also investigated the effect of the Mhp V273A mutation on the formation of the ThsB1/B2 complex. To do this we performed pull-down experiments in staphylococci harboring pMhp plasmids for the over-expression of wild-type and V273A mutant Mhp, in the absence of phage infection. Similarly to the results obtained during induction of the ΦNM1 prophage, SDS-PAGE of the proteins copurified with hexa-histidyl versions of ThsB1 or ThsB2, indicated the presence of a complex composed of both proteins and wild-type Mhp (Fig. 3C). This tripartite complex, however, was not captured during expression of Mhp^V273A^. Instead, we observed that the mutant Mhp co-purified with ThsB1, while ThsB2 did not interact with ThsB1 nor Mhp^V273A^ (Fig. 3C). This result demonstrates that the V273A escaper mutation prevents the formation of the ThsB1/B2/Mhp complex.

**Figure 3.**
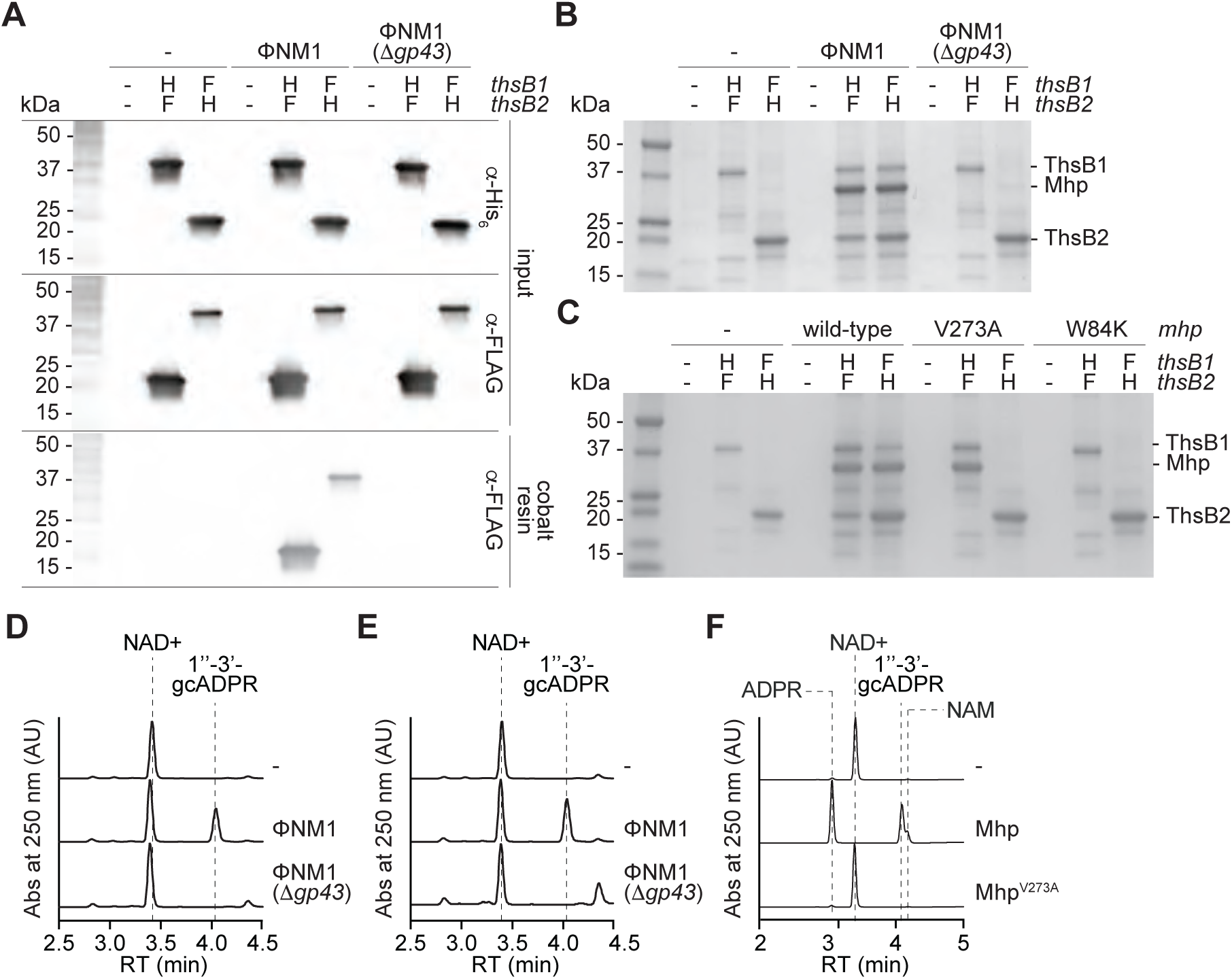
ThsB1/B2 form a complex with the major head protein *in vivo*. **(A)** Immunoblot analysis of proteins extracted from staphylococci expressing hexahystidyl-(H) or FLAG-(F) tagged versions of ThsB1 or ThsB2, uninfected or infected with wild-type or Δ*gp43* ΦNM1 phage, either before (input) of after affinity chromatography using a cobalt resin. Proteins were separated by SDS-PAGE, and electrotransferred to a PVDF membrane. Tagged proteins were detected with anti-hexahystidyl (α-His) or anti-FLAG (α-FLAG) antibodies and chemiluminescence staining. **(B)** Coomassie Blue-stained SDS-PAGE of proteins isolated after cobalt resin affinity chromatography in the experiment described in **(A)**. His_6_-ThsB1, 41.8 kDa; ThsB2-His_6_, 23.4 kDa; Mhp, 36.8 kDa. Protein molecular weight (kDa) markers are shown. **(C)** Coomassie Blue-stained SDS-PAGE of proteins isolated from staphylococci expressing hexahystidyl-(H) or FLAG-(F) tagged versions of ThsB1 and ThsB2 and either wild-type, V273A or W84K Mhp, in the absence of phage infection, after cobalt resin affinity chromatography. Protein molecular weight (kDa) markers are shown. **(D)** HPLC analysis of the products resulting from the incubation of the proteins purified from staphylococci expressing His_6_-ThsB1 and ThsB2-FLAG, uninfected or infected with wild-type or Δ*gp43* ΦNM1 phage, with NAD+, using a cobalt resin. Retention times (RT) of reactants and products are marked by dotted lines. **(E)** Same as **(D)** but using proteins extracted from staphylococci expressing ThsB1-FLAG and ThsB2-His_6_. **(F)** Same as **(D)** but adding purified His_6_-ThsA to the reaction.

Finally, we determined whether the isolated complexes were catalytically active. First, the complex purified after prophage induction (Figs. 3A-B) was incubated with NAD+ to test their cyclase activity (Fig. S3C) using high-performance liquid chromatography (HPLC) to detect the generation of gcADPR. To be able to interpret the resulting chromatograms, we determined the retention times for NAD+ and both possible cyclic products, 1’’-2-gcADPR and 1’’-3-gcADPR (Fig. S3D). We found that the complex pulled-down by His_6_-ThsB1 or ThsB2-His_6_ produced 1”-3’-gcADPR only when the wild-type prophage was induced; not in the absence of a prophage or during induction of ΦNM1(11*gp43*) (Fig. 3D-E). To test for the full Sau-Thoeris response, we determined whether 1”-3’-gcADPR produced by the ThsB1/B2/Mhp complex can activate ThsA to cleave NAD+ into nicotinamide (NAM) and ADPR (Fig. S3E). The complexes pulled-down by His_6_-ThsB1 in the presence of ThsB2-FLAG and wild-type or V273A mutant major head protein (Fig. 3C) were incubated with NAD+ and purified His_6_-ThsA. After establishing the retention times of the cleavage products (Fig. S3D; note that NAM absorbance at 250 nm is much lower than that of the other compounds used in this study), we found that the complex enriched in the presence of wild-type Mhp, but not Mhp^V273A^, were able to stimulate ThsA to produce NAM and ADPR (Fig. 3F). Based on these results, we conclude that Sau-Thoeris immunity is initiated by the interaction between viral Mhp, ThsB1 and ThsB2, a tripartite complex that cannot form during infection with escaper phages expressing Mhp^V273A^.

### ThsB1 interacts with Mhp to recruit ThsB2 and stimulate its cyclase activity

The presence of two ThsB subunits, both essential for immunity (Fig. 1E), is an intriguing feature of the Sau-Thoeris system; their individual functions and how they coordinate them for the synthesis of gcADPR during phage infection is not known. The results presented in Figure 3C demonstrated that ThsB1 and ThsB2 do not interact with each other in the absence of Mhp. In addition, the experiment suggested a possible scenario in which, during infection, the interaction between Mhp and ThsB1 recruits ThsB2 to generate an active complex capable of catalyzing the cyclization of NAD into gcADPR, and that the V273A mutation prevents ThsB2 recruitment. We tested this model by expressing hexa-histidyl-tagged versions of ThsB1 or ThsB2 alone, not as a pair, infecting the cultures with Φ80α-vir (MOI 10) for 20 minutes and capturing each sensor protein using cobalt affinity chromatography. SDS-PAGE of the purified proteins showed that Mhp co-purifies with ThsB1, but not ThsB2 (Fig. 4A), a result that supports our model of Mhp activation.

**Figure 4.**
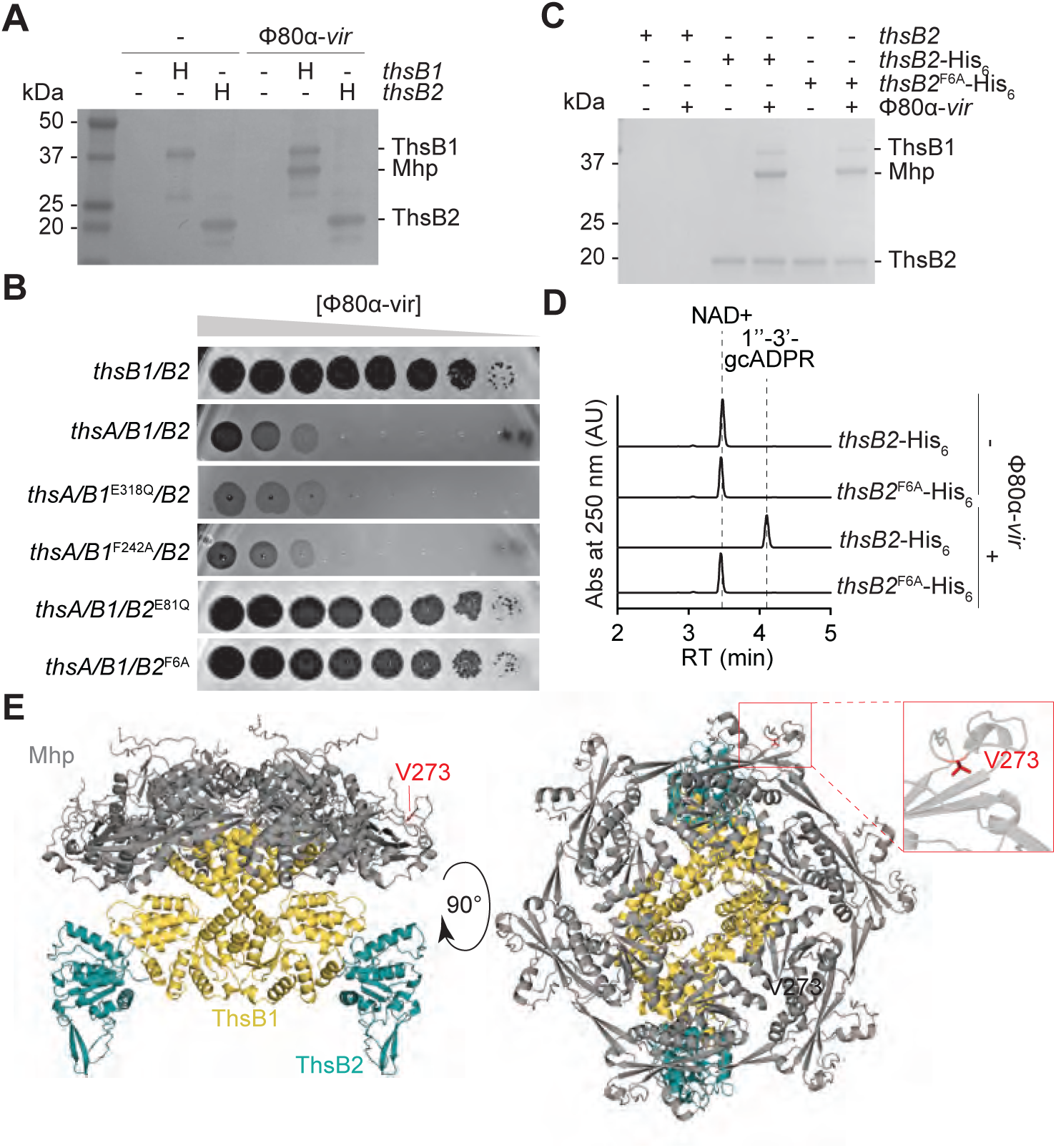
ThsB1 interacts with Mhp to recruit ThsB2 and stimulate its cyclase activity. **(A)** Coomassie Blue-stained SDS-PAGE of proteins isolated from staphylococci expressing hexahystidyl-(H) tagged versions of ThsB1 or ThsB2, uninfected or infected with Φ80α-vir, after cobalt resin affinity chromatography. Protein molecular weight (kDa) markers are shown. **(B)** Tenfold serial dilutions of Φ80α-vir on lawns of *S. aureus* RN4220 harboring plasmids carrying either an incomplete (*thsB1/B2*) or full (*thsA/B1/B2*) Thoeris operon carrying wild-type or mutant versions of *thsB1* or *thsB2*. **(C)** Coomassie Blue-stained SDS-PAGE of proteins isolated from staphylococci expressing ThsB2, ThsB2-His_6_ or ThsB2^F6A^-His_6_, uninfected or infected with Φ80α-vir, after cobalt resin affinity chromatography. Protein molecular weight (kDa) markers are shown. **(D)** HPLC analysis of the products resulting from the incubation of the proteins purified from staphylococci expressing ThsB2-His_6_ or ThsB2^F6A^-His_6_, uninfected or infected with Φ80α-vir, with NAD+, using a cobalt resin. Retention times (RT) of reactants and products are marked by dotted lines. **(E)** AlphaFold3 structure of a complex formed by Φ80α Mhp (hexamer; grey) ThsB1 (two copies; yellow) and ThsB2 (two copies; teal). Two angles of the structure (90° rotation), as well as the position of residue V273 (red) are shown.

Next, we investigated which subunit performs the cyclase reaction. To do this, we generated AlphaFold structures for ThsB1 (Fig. S4A) and ThsB2 (Fig. S4B) and aligned them to closely related and previously characterized TIR proteins *Bacillus cereus* ThsB (BcThsB; PDB: 6LHY), *Acinetobacter baumannii* TIR domain (AbTir; PDB: 7UXU), human sterile alpha and TIR motif containing preotein 1 (SARM1; PDB: 6O0R) and *Arabidopsis thaliana* resistance protein RPP1 (PDB: 7DFV) (Fig. S4C). This comparison revealed a conserved active site featuring the catalytic glutamate located across a phenylalanine residue (which in other related proteins could be either an alanine or a tyrosine) (Shi et al., 2024), and enabled us to identify the putative active sites for ThsB1 (containing E318 and F242; Fig. S4C) and ThsB2 (containing E81 and F6; Fig. S4C). We made double substitutions of glutamate for glutamine and phenylalanine for alanine in both proteins and found that the ThsB2 mutations (E81Q and F6A), but not the ThsB1 mutations (E318Q and F242A), disrupted immunity against Φ80α-vir in a plaquing assay (Fig. 4B). These results suggest that the catalytic activity of ThsB2 is required for the synthesis of the second messenger. To test this, we expressed ThsB2^F6A^-His_6_ along with ThsB1 and ThsA, and performed pulldown experiments using lysates of staphylococci infected with Φ80α-vir at an MOI 10, collected 20 minutes post-infection. We found that the F6A mutation does not interfere with the formation of the complex with ThsB1 and Mhp (Fig. 4C). We then tested the isolated complex for cyclase activity *in vitro*. As opposed to the complex isolated after pull-down of wild-type ThsB2-His_6_, we were unable to detect the generation of gcADPR, even in the presence of a wild-type ThsB1 (Fig. 4D). Altogether, these results support a model in which, upon phage infection, viral Mhp and ThsB1 interact to recruit ThsB2 and stimulate its NAD+ cyclase activity.

### Sau-Thoeris senses the hexameric form of Mhp

The structure of Mhp from Φ80α [Mhp(Φ80α)] has been solved experimentally (Fig. S5A) and demonstrated to be a hexamer (Dearborn et al., 2017) (Fig. S5B). Importantly, oligomeric complexes of the major head protein can form in the absence of other structural components of the procapsid (Spilman *et al*., 2011). We used AlphaFold3 to explore possible interactions between this structure and ThsB1 and ThsB2. Supporting the results showing co-purification between Mhp and ThsB1, but not ThsB2 (Fig. 4A), the predicted structure showed that ThsB1 forms a dimer that bridges Mhp to the ThsB2 cyclase, directly interacting with the center of the Mhp hexameric ring on one side, and with two ThsB2 monomers on the other (Fig. 4E). In this AlphaFold3 model, V273 is located in the perimeter of the ring and does not make a direct contact with ThsB1 (Fig. 4E, inset), a prediction consistent with the experimental result showing that the V273A escape mutation does not prevent the interaction of ThsB1 and Mhp (Fig. 3C). In addition, the structural model of the mutant hexamer differs from that formed by wild-type Mhp (Fig. S5C; root mean square deviation of atomic positions (RMSD) values 10.698 Å, 12,922 atoms). Therefore, the combination of the different AlphaFold3 predictions with the available escaper mutant experimental data altogether suggest that ThsB1 senses the hexameric complex formed by Mhp.

The Mhp complex is held together through the interaction of two loops, the 12-residue P-loop and the 30-residue E-loop, located approximately 60 Å apart at opposite ends of the monomer (Johnson and Chiu, 2007; Spilman *et al*., 2011) (Fig. S5A). E- and P-loops from different monomers associate with each other to form the hexameric ring of Mhp (inset, Fig. S5B). V273 is situated within the P-loop and therefore we believe that the mutation to alanine could affect the interaction with the E-loop to generate an altered hexameric conformation (Fig. S5C). We wondered whether mutations in the E-loop could result in similar structural variations of the capsid complex that would prevent the activation of the ThsB1/B2 complex. We used the ConSurf database (Ben Chorin et al., 2020) to look for conserved residues within the E-loop, which allowed us to identify a tryptophan residue in position 84, which is highly conserved but substituted by a lysine in some sequences. We speculated that the mutation W84K would be tolerated and lead to the formation of a functional capsid that, given the divergent chemical properties of these two amino acids, could display an altered conformation, and possibly escape ThsB1 recognition. We first checked that the mutation did not disrupt capsid formation. We induced a ΦNM1(Δ*gp43*) lysogen harboring an empty vector control or a plasmid expressing either wild-type Mhp, Mhp^V273A^ or Mhp^W84K^ from an IPTG-inducible promoter. After induction of the defective prophage with MMC, Mhp^W84K^ enabled the formation of as many PFUs as both wild-type and V273A mutant capsid proteins (Fig. S5D). We also engineered this mutation into Φ80α-vir and found that it enabled similar levels of escape from Sau-Thoeris immunity as the V273A mutation (Fig. S2C). We then assessed whether over-expression of Mhp^W84K^ from a plasmid was sufficient to activate the Sau-Thoeris response, and found that the mutant Mhp was unable to mediate the reduction of NAD+ levels observed after over-expression of the wild-type capsid protein (Fig. 2D). Similarly to the AlphaFold3 model of hexameric Mhp^V273A^, the prediction for the Mhp^W84K^ mutant showed alterations in the capsid complex that could explain the inability to stimulate ThsB1/B2 cyclase activity (Fig. S5E; RMSD=11.970 Å, 13,233 atoms). We then performed pull-downs of His_6_-ThsB1 or ThsB2-His_6_ in the presence of FLAG-tagged ThsB2 or ThsB1, respectively, after over-expression of Mhp^W84K^, and found that the mutant capsid protein did not interact with either ThsB1 or ThsB2 (Fig. 3C). This result demonstrated that the W84K substitution prevents the formation of the Mhp/ThsB1/ThsB2 complex even in the presence of an intact P-loop. Given that V273 and W84 are far apart in the Mhp monomer but located closely in the hexameric form of this protein, the finding that mutations in either of these residues prevents Sau-Thoeris activation strongly suggests that the association between capsid proteins, and not the Mhp monomer, is required for the sensing of phage infection by ThsB1/B2.

### Viral major head protein is sufficient to stimulate ThsB1/B2 cyclase activity *in vitro*

Given that the experiments described above involved the pull-down of proteins from full cell extracts, they cannot rule out whether other cell components not detected by SDS-PAGE, in addition to Mhp, are necessary for the stimulation of ThsB1/B2 cyclase activity. Therefore, we investigated the minimal requirements for ThsB1/B2 activation by performing biochemical reactions with purified proteins and analyzing the reaction products using HPLC. We obtained pure preparations of His_6_-ThsA, His_6_-ThsB1, and ThsB2-His_6_ through affinity chromatography of staphylococcal cell lysates over-expressing the tagged proteins using cobalt resin (Fig. 5A). Because Mhp did not tolerate the addition of affinity tags, we used a method previously developed (Spilman *et al*., 2011) to purify Mhp and the V273A mutant, from lysates of *S. aureus* RN4220 over-expressing these proteins (harboring pMhp plasmids, see above), using PEG precipitation followed by separation in a sucrose gradient (Fig. 5B). We first tested whether His_6_-ThsB1 and/or ThsB2-His_6_ possess NAD+ cyclase activity (Fig. S3C) in the presence of wild-type or V273A mutant Mhp. We found that wild-type Mhp only stimulates the production of 1’’-3’ gcADPR when both ThsB proteins are present (Fig. 5C), but that Mhp^V273A^ was incapable of triggering the cyclase reaction (Fig. 5D). Next, we corroborated the ability of purified His_6_-ThsA to cleave NAD+ into ADPR and NAM (Fig. S3E) (Ofir *et al*., 2021; Tamulaitiene *et al*., 2024) in the presence of commercially available 1’’-3’ gcADPR (BioLog) (Fig. 5E). Finally, we recapitulated the full Sau-Thoeris response *in vitro* by mixing the His_6_-ThsB1 and ThsB2-His_6_ sensor cyclase, the His_6_-ThsA effector NADase, the NAD+ substrate for both reactions, and the Mhp or Mhp^V273A^ activators. We found that only the wild-type activator promoted the conversion of NAD+ into ADPR and NAM (Fig. 5F). Altogether, these data demonstrate that ThsB1 and ThsB2 are activated by Mhp to synthesize 1’’-3’ gcADPR, which in turn activates ThsA.

**Figure 5.**
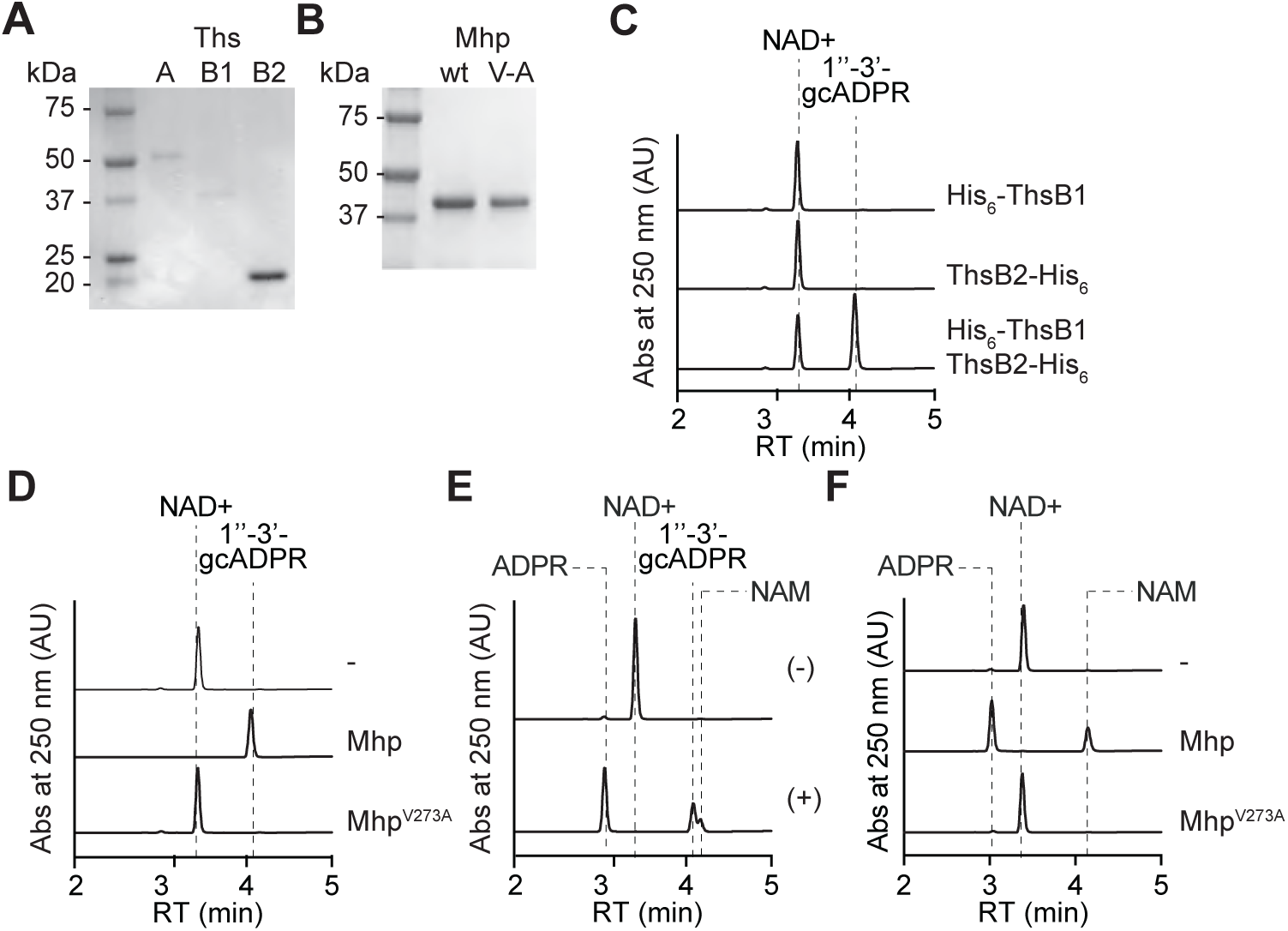
Viral major head protein is sufficient to stimulate ThsB1/B2 cyclase activity *in vitro*. **(A)** Coomassie Blue-stained SDS-PAGE of proteins purified from *E. coli* expressing His_6_-ThsA (56.0 kDa), His_6_-ThsB1 (41.8 kDa), and ThsB2-His_6_ (23.4 kDa) using a cobalt resin. Protein molecular weight (kDa) markers are shown. **(B)** Coomassie Blue-stained SDS-PAGE of Mhp proteins, wild-type and V273A mutant (V/A) purified from staphylococci harboring pMhp plasmids, using PEG-enrichment. Protein molecular weight (kDa) markers are shown. **(C)** HPLC analysis of the products resulting from the incubation of purified ThsB2-His_6_, ThsB2^F6A^-His_6_ or both, with NAD+. Retention times (RT) of reactants and products are marked by dotted lines. **(D)** Same as **(C)** but after incubation of both ThsB2-His_6_ and ThsB2^F6A^-His_6_, alone (-) or in the presence of purified wild-type or V273A mutant Mhp. **(E)** HPLC analysis of the products resulting from the incubation of purified His_6_-ThsA and commercially available 1’’-3’ gcADPR. Retention times (RT) of reactants and products are marked by dotted lines. **(F)** Same as **(D)** but in the presence of purified His_6_-ThsA.

### ThsB1/2 recognize conserved capsid proteins of diverse staphylococcal phages

The above results demonstrated that the Mhp from the staphylococcal phages Φ80α-vir and ΦNM1γ6 interact with ThsB1 to activate the Sau-Thoeris response. Our initial experiments indicated that this system also provides immunity against ΦNM4γ4, ΦJ1, ΦJ2, ΦJ4, but not Φ12γ3 (Fig. 1B). Therefore, we hypothesized that ThsB1/B2 would recognize the major head protein of these phages, except for that of Φ12γ3, to trigger the Sau-Thoeris response. To investigate whether the sequence and/or structure of Mhp(Φ12γ3) is fundamentally different than that of the Mhps derived from the rest of the phages tested in this study, we performed a multiple sequence alignment to generate the phylogenetic tree of the six Mhps, using the ΦNM1γ6 homolog as the reference sequence. Indeed, we found that Mhp(Φ12γ3) is the most distant member of this group of structural proteins (Fig. S6A). Sequence divergence was reflected in a marked structural variation (based on AlphaFold3 predictions) for Mhp(Φ12γ3). We quantified these differences by comparing the structure of each predicted Mhp monomer to that of Mhp(ΦNM1γ6) and calculated the RMSD values. While Mhps from phages Φ80α-vir, ΦNM4γ4, ΦJ1/2, ΦJ4 display a very similar folding to Mhp(ΦNM1γ6), with RMSD values around 1 Å, the RMSD value of the predicted structure of Mhp(Φ12γ3) was over 27 Å (Fig. S6B). The structural disparity of the Mhp(Φ12γ3) monomer is also reflected in the predicted hexameric ring formed by this protein, which is notably to the hexamers predicted for the rest of the Mhps investigated in this study (Fig. S6C).

To test whether the sequence and structural predictions correlate with the ability of the different Mhps to activate the Sau-Thoeris response, we cloned the major head protein gene of all these phages (ΦJ1 and ΦJ2 Mhp have the same amino acid sequence) and tested their effect on different assays previously used to characterize the activating properties of Φ80α and ΦNM1 Mhps. We first investigated whether overexpression of Mhp was sufficient to stimulate Sau-Thoeris in the absence of infection, *in vivo*. Indeed, Mhp from all phages but Φ12γ3 was sufficient to cause cell arrest in the presence of ThsA/B1/B2/A, but not ThsB1/B2 alone, measured as the OD_600_ of the induced cultures (Fig 6A and Fig. S6D), as well NAD+ depletion (Fig. 6B). We also purified Mhp from staphylococci infected with these phages (Fig. S6E) and incubated them with purified His_6_-ThsB1, ThsB2-His_6_ and NAD+ to test for their ability to stimulate gcADPR production *in vitro.* Similarly to the *in vivo* results, we found that Mhp from ΦNM4γ4, ΦJ1/2, and ΦJ4, but not Φ12γ3, activated the synthesis of 1’’-3’ gcADPR (Fig. 6C); which in turn was able to induce the cleavage of NAD+ when mixed with purified His_6_-ThsA (Fig. S6F). These results demonstrate that diverse staphylococcal phages have structurally conserved capsid proteins that are recognized by Sau-Thoeris to trigger defense.

**Figure 6.**
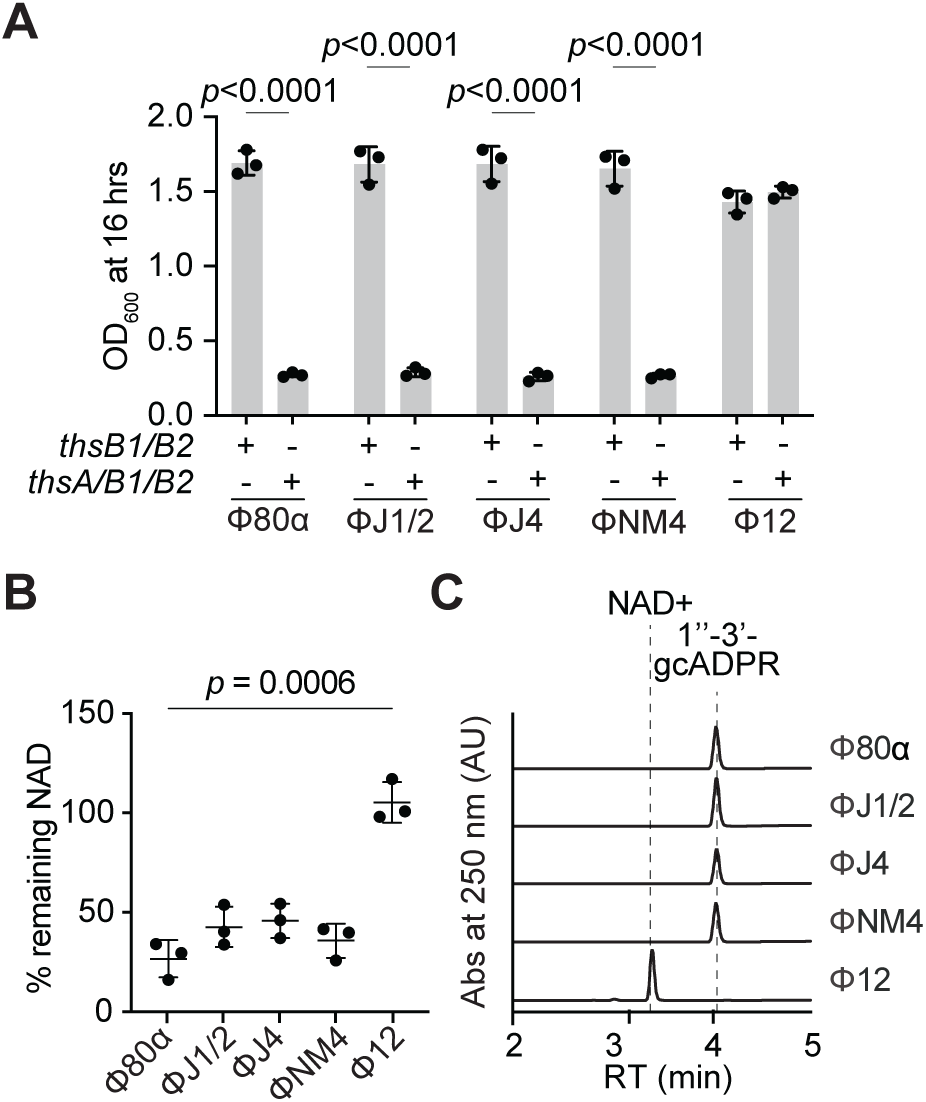
ThsB1/2 recognize conserved capsid proteins of diverse staphylococcal phages. **(A)** Growth of *S. aureus* RN4220 harboring plasmids carrying either an incomplete (*thsB1/B2*) or full (*thsA/B1/B2*) Thoeris operon in the presence of a second plasmid expressing Mhp from different staphylococcal phages, determined as the OD_600_ of the cultures 16 hours after addition of IPTG. Mean of +/- S.D. of three biological replicates is reported; *p* value was obtained using an unpaired, two-tailed, *t*-test. **(B)** Measure of % remaining NAD+/NADH (NAD), calculated as the ratio of the concentration of NAD+ and NADH detected in staphylococci harboring a plasmid carrying a full (*thsA/B1/B2*) Thoeris operon, to the value detected in the presence of an incomplete (*thsB1/B2*) system, after induction of the expression of Mhp from different staphylococcal phages. Mean of +/- S.D. of three biological replicates is reported; *p* value was obtained using an unpaired, two-tailed, *t*-test**. (C)** HPLC analysis of the products resulting from the incubation of purified ThsB2-His_6_ and ThsB2^F6A^-His_6_ with NAD+, in the presence of purified Mhp from different staphylococcal phages. Retention times (RT) of reactants and products are marked by dotted lines.

## DISCUSSION

Here we investigated how the *Staphylococcus aureus* Thoeris system is activated by phage during infection. We found that the expression of the major head proteins of different staphylococcal Siphoviridae phages that are susceptible to Thoeris defense mediates the association of the TIR-containing proteins ThsB1 and ThsB2. Structural predictions as well as experimental data support a model in which binding of ThsB1 to the hexameric complex formed by the capsid proteins leads to the recruitment of ThsB2 and stimulation of its cyclase activity, which converts NAD+ into the second messenger 1’’-3’-gcADPR. The results that validate this mechanism of Sau-Thoeris activation are: (i) ThsB1 and ThsB2 do not interact with each other in the absence of Mhp (Fig. 3A), (ii) Mhp expression leads to its association with both ThsB1 and ThsB2 (Fig. 3B), (iii) in the absence of ThsB2, ThsB1 interacts with Mhp, but ThsB2 does not interact with Mhp in the absence of ThsB1 (Fig. 4A), and (iv) the ThsB2 putative cyclase active site is critical for the synthesis of gcADPR. An AlphaFold3 prediction for the structure of a complex formed by Mhp, ThsB1 and ThsB2 independently aligned with our experimental data, as it showed a ThsB1 dimer interacting with a hexameric Mhp capsid complex on one side of the dimer and with two individual ThsB2 subunits on the other side (Fig. 4E).

There are several features of this model that will require further investigation. For example, how ThsB1 binds the Mhp hexamer is not completely clear. Our data indicates that mutations in the P-loop (V273A) or E-loop (W84K) prevent Sau-Thoeris activation. Because these loops are 60 Å apart in the Mhp monomer but overlap in the Mhp hexamer, we conclude that ThsB1 must recognize the hexameric conformation of this phage protein (ThsB1 would have to bind across the full length of the Mhp monomer to be able to interact with both ends of the protein). However, it remains possible that ThsB1 associates with the vertex of the Mhp hexamer, where the overlapping loops that hold together the capsid complex are located, and not with the center of the ring as predicted by AlphaFold3. In this case the V273A and W84K escape mutations would directly affect ThsB1 role in Sau-Thoeris activation, instead of affecting the hexameric conformation of Mhp, as we propose. Finally, the changes experienced by ThsB2 as it goes from an inactive monomer to an active cyclase in association with ThsB1 and Mhp, remain to be determined. We believe that future structural studies of the Mhp/ThsB1/ThsB2 and Mhp/ThsB1 complexes, in the presence and absence of the NAD+ substrate, will clarify these aspects of our model.

An interesting finding from our work is that bacterial TIR proteins can cooperate to provide defense. While the best characterized Thoeris defense system, from *Bacillus spp.*, possess a single ThsB subunit, those that encode two ThsB units have been shown to employ each TIR protein to independently sense different phages (Ofir *et al*., 2021). Although we cannot rule out the possibility that ThsB1 or ThsB2 are directly activated by phages not used in this study, our data shows, at least for the staphylococcal Siphoviridae phages we tested, an interplay between bacterial TIR proteins that is somewhat reminiscent of the interaction between TIR proteins in eukaryotes (Wang et al., 2006). For example, the TIR proteins TLR1 and TLR2, located in the plasma membrane, interact with each other to sense ligands derived from bacterial envelopes. Another example is the pairwise cooperation of TLR8 with TLR7 or TLR9 within endosomal membranes to regulate the eukaryotic inflammatory response upon detection of viral nucleic acids produced during infection (Wang *et al*., 2006).

There are other characterized defense strategies that recognize capsid proteins as a sign of infection. In *Escherichia coli*, the major capsid protein of phage SECΦ27 and other related phages, directly activates the CapRel toxin-antitoxin system commonly present in prophages (Zhang et al., 2022). CapRel is a single polypeptide folding into a “closed” conformation in which the C-terminal domain prevents the toxic activity of the N-terminal domain. Direct binding of the viral capsid protein to the inhibitory domain of CapRel releases the toxic domain, triggering abortive infection immunity. Also in *E. coli*, the Lit protease that causes translation inhibition during T4 infection, can be activated by a peptide derived from the viral capsid protein Gp23, in the absence of phage (Bergsland et al., 1990). *E. coli* CBASS systems can be activated by phage capsid proteins as well. The prohead protease of phage BAS13 interacts and stimulates the type I CBASS cyclase EcCdnD12 and it is sufficient to induce cells death in a CBASS-dependent manner in vivo (Richmond-Buccola et al., 2024). In addition, mutations in capsid-encoding genes of *Pseudomonas* and *Staphylococcus* phages have been found to avoid CBASS immunity (Banh *et al*., 2023; Huiting et al., 2023). Finally, phage T5 accumulates mutations in the major capsid protein precursor pb8 to evade Pycsar immunity (Tal et al., 2021) and phage T7 escapes F restriction in *E. coli* through mutations in major capsid protein gene 10 (Molineux et al., 1989). These findings suggest an involvement of phage capsids in the activation of prokaryotic immunity. In eukaryotes, TIR protein-based immunity can also be triggered by the recognition of viral structural components. This is the case for TLR2, present in the surface of primary human liver cells, which is activated by the adeno-associated virus capsids to induce the production of inflammatory cytokines (Hosel et al., 2012). Therefore, our results demonstrate a conserved mechanism for the recognition of viral structural components by TIR-containing proteins to start innate immunity against infection.

Immunological logic dictates that bacterial defense systems sense conserved molecules produced during phage infection (Gao et al., 2022; Stokar-Avihail et al., 2023). Conservation of immunological targets ensures (i) that the activating molecules are present in many viruses, making the immune system useful against a broad range of phages, and (ii) that the target is essential for optimal phage propagation and therefore difficult to mutate, reducing the chances of viral escape. We believe that the targeting of Mhp by Sau-Thoeris (and of other capsid proteins by other defense systems) meets both evolutionary requirements. Mhp from Φ80α is highly conserved among many staphylococcal phages (Fig. S6A) and four out of five homologs were able to activate Sau-Thoeris (Figs. 6 and S6D). Mhp is also essential for Φ80α propagation (Fig. S2C) and we were able to find only two escape mutations, V273A and W84K, using either sequential infections staphylococci carrying the Sau-Thoeris system or genetic engineering based on sequence conservation, respectively. Mhp mutations that escape Sau-Thoeris cannot disrupt hexamer formation, but instead result in the generation of an altered hexameric conformation that cannot activate ThsB1/ThsB2 but can still assemble into procapsids. Although such mutations are infrequent, they can accumulate as a consequence of the evolutionary arms race between phages and their prokaryotic hosts, and most likely will result in changes in capsid morphology. All the phages in this study belong to the Siphoviridae group, which can be classified into distinct serogroups based on morphological differences in their head structures (Xia and Wolz, 2014).

Notably, phages Φ80α, ΦNM1, ΦNM4, ΦJ1, ΦJ2, and ΦJ4, which activate Sau-Thoeris, belong to serogroup B and have isometric capsids. In contrast, Φ12, whose Mhp avoids triggering TIR-mediated defense, belongs to serogroup A and has a more prolate head structure, lengthened in one direction (Xia and Wolz, 2014). We believe that the evasion of Sau-Thoeris immunity, and possibly other defense systems that are activated by capsid proteins, represents one important evolutionary force in the differentiation of the Φ12 capsid structure, and, more generally, in the generation of structural diversity in staphylococcal phages.

## METHODS

### Bacterial strains and growth conditions

The bacterial strains used in this study are listed in Supplementary Methods Table 1. *Staphylococcus aureus* strain RN4220 (Xia and Wolz, 2014) was grown at 37°C with shaking (220 RPM) in brain heart infusion (BHI) broth, supplemented with chloramphenicol (10 μg mL^-1^) or erythromycin (10 μg mL^-1^) to maintain pC194-based (Horinouchi and Weisblum, 1982a) or pE194-based plasmids (Horinouchi and Weisblum, 1982b), respectively. Cultures were supplemented with erythromycin (5 μg mL^-1^) to select for strains with chromosomally integrated Sau-*Thoeris* or Sau-*thsB1/B2*. Gene expression was induced by the addition of 1 mM isopropyl-d-1-thiogalactopyranoside (IPTG), where appropriate.

### Bacteriophage propagation

The bacteriophages used in this study are listed in Supplementary Methods Table 2. To generate a high titer phage stock, an overnight culture of *S. aureus* RN4220 was diluted 1:100 and outgrown to mid-log phase (∼90 min) in BHI broth supplemented with 5 mM CaCl_2_. The culture was diluted to an optical density measurement at 600 nm (OD_600_) of 0.5 (∼1×10^8^ CFU mL^-1^). The culture was infected by adding phage at a multiplicity of infection (MOI) of 0.1 (∼1×10^7^ PFU mL^-1^), or by inoculating with either a single picked plaque or scrape of a frozen stock. The infected culture was grown at 37°C with shaking and monitored for lysis (full loss of turbidity was typically observed ∼3-4 hr). Culture lysates were centrifugated (4,300 x g for 10 min) to pellet cellular debris. The supernatant was collected, passed through a sterile membrane filter (0.45 μm), and stored at 4°C. Phage concentrations were determined by serially diluting the obtained stock in 10-fold increments and spotting 2.5 μL of each dilution on BHI soft agar mixed with RN4220 and supplemented with 5 mM CaCl_2_. After incubation overnight at 37°C, individual plaques (i.e. zones of no bacterial growth) were counted, and the viral titer was calculated.

### Molecular cloning

The plasmids (and details of their construction) and the oligonucleotide primers used in this study are listed in Supplementary Methods Table 3 and Supplementary Methods Table 4, respectively. The coding sequences of Sau-Thoeris and phage gene products were obtained from G blocks, genomic DNA preparations or phage stocks, respectively.

### Chromosomal integration of Sau-Thoeris

Sau-Thoeris or Sau-ThsB1/B2, along with an erythromycin resistance (ermR) cassette, was integrated into the *hsdR* gene (which encodes the defective R-subunit of the restriction-modification system in *S. aureus* RN4220), an insertion site which was previously shown to not impact growth (Maguin et al., 2022). *Sau-thsA/B1/B2-ermR* and *Sau-thsB1/B2-ermR* were amplified from the plasmids pDVB223 and pCF11 respectively, using primers oCR482 and oCR483 or oCR484, which were flanked with loxP sites at both ends followed by 60-bp homology regions to *hsdR*. Electrocompetent *S. aureus* RN4220 cells harboring the recombineering plasmid pPM300 (Banh *et al*., 2023) were electroporated with 1-2 μg of PCR product and selected for with erythromycin (5 μg mL^-1^). Potential integrants were screened by colony PCR as well as for functional immunity, and then verified by Sanger sequencing.

### Prophage recombineering

The prophage strains and the oligonucleotide primers used in this study are listed in Supplementary Methods Tables 1 and 4. A chloramphenicol resistance (cmR) cassette flanked by loxP sites and 60 bp homology regions, were integrated within codons for phage genes of interest corresponding to the homology overhangs. The loxP CmR was amplified from a G-block (Azenta), using primers oCR24 and oCR25, oCR26 and oCR37, oCR30 and oCR31, oCR32 and oCR33, oCR485 and oCR486, oCR487 and oCR488, oCR489 and oCR490, oCR491 and oCR492, oCR493 and oCR494, oCR495 and oCR496, which were all flanked with 60-bp homology regions. Electrocompetent *S. aureus* RN4220 cells harboring the recombineering plasmid pPM300 were electroporated with 1-2 μg of PCR product and selected for with chloramphenicol (5 μg mL^-1^). Potential integrants were screened by colony PCR as well as for functional immunity, and then verified by Sanger sequencing.

### Generation of Φ80α-vir(*mhp^W84K^)*

Wild-type Φ80α-vir was passaged on a liquid culture of *S. aureus* RN4220 harboring a plasmid (pCR186) encoding the *mhp^W84K^* gene flanked by 500-nt upstream and downstream homology arms corresponding to Φ80α *gp46* and *gp48*, respectively. To isolate individual plaques, the lysed culture supernatant spotted onto a lawn of RN4220 harboring a type II-A Sau CRISPR-Cas targeting plasmid (pCR187) in BHI soft agar for counter-selection against wild-type phage and enrichment of Φ80α-vir::*mhp^W84K^*. The mutation was confirmed by Sanger sequencing.

### Soft-agar phage infection

100 μL of an overnight bacterial culture was mixed with 5 mL BHI soft agar supplemented with 5 mM CaCl_2_ and poured onto BHI agar plates to solidify at room temperature (∼15 min). Phage lysates were serially diluted 10-fold and 2.5 μL was spotted onto the soft agar surface. Once dry, plates were incubated at 37°C overnight and visualized the next day. Individual plaques (zones of no bacterial growth) were enumerated manually.

### Liquid culture phage infection

Overnight cultures were diluted 1:100 in BHI supplemented with 5 mM CaCl_2_ and the appropriate antibiotic for selection, outgrown at 37°C with shaking to mid-log phase (∼90 min), and normalized to OD_600_ 0.5. For the desired MOI, a calculated volume of phage stock was added to each culture and 150 μL was seeded into each well of a 96-well plate. OD_600_ was measured every 10 min in a microplate reader (TECAN Infinite 200 PRO) at 37°C with shaking.

### Protein expression and purification

ThsA, ThsB1, and ThsB2 were expressed and purified using the following approach: transformed *S. aureus* RN4220 were grown in BHI broth with 1 mM IPTG at 37°C with shaking to OD_600_ 1, at which point the culture was cooled on ice for 10 min and bacteria were harvested. Pellets were resuspended in lysis buffer (25 mM Tris pH 7.4, 100 mM NaCl, 10% glycerol, 2 mM *β*-mercaptoethanol, 5 mM MgSO_4_), and subjected to a single freeze-thaw cycle. The cells were incubated at 37°C with Lysostaphin, DNase I, and EDTA-free protease inhibitor cocktail for 30 min. After incubating, the cells were lysed using sonication (70% amplitude, 10 sec on/off, 2 min total). Lysates were clarified by centrifugation and applied to cobalt affinity resin. After binding, the resin was washed extensively with high salt lysis buffer (500 mM) prior to elution with lysis buffer containing 200 mM imidazole. Eluted proteins were subjected to overnight 4°C dialysis into reaction buffer (25 mM Tris pH 7.4, 100 mM NaCl, 10% glycerol, 2 mM *β*-mercaptoethanol). The next day, proteins were concentrated using 10,000 MWCO centrifugal filters (Amicon). Purified proteins were visualized by SDS-PAGE and used for downstream *in vitro* assays.

### Purification of native major head proteins

Native major head proteins from Φ80α, ΦNM1, ΦNM4, ΦJ1, ΦJ2, ΦJ4, and Φ12 were expressed and purified either according to an established protocol (Johnson and Chiu, 2007; Spilman *et al*., 2011) using *S. aureus* cells harboring pMhp or as follows: PEG-precipitated phage particles were resuspended in unfolding buffer (4M guanidine-HCL, 50 mM Tris-HCl pH 8.0, and 150 mM NaCl). Proteins were then layered on a 10-40% sucrose gradient and separated by centrifugation at 100,000g for 2 hours. The gradients were manually fractionated from the top and each fraction was analyzed by SDS-PAGE. Fractions with capsid protein were subjected to dialysis into refolding buffer (25 mM Tris pH 7.4, 100 mM NaCl, 10% glycerol, 2 mM *β*-mercaptoethanol, 5 mM MgSO_4_) overnight at 4°C. The final precipitate was removed by centrifugation and the supernatant was used for downstream enzymatic assays.

### Nucleotide synthesis assays

Nucleotide synthesis assays were performed using a variation of the method described by (Ka *et al*., 2020). The final reactions (25 mM Tris pH 7.4, 100 mM NaCl, 10% glycerol, 2 mM *β*-mercaptoethanol, 100 *μ*M NAD+, 1 uM head protein, and 1 μM enzyme) were started with the addition of enzyme. All reactions were incubated for 2 hours at 37°C. To isolate the gcADPR product for HPLC analysis, nucleotide synthesis reaction conditions were scaled up to 200 *μ*L reactions. Reactions were incubated with gentle shaking for 2 hr at 37°C. Following incubation, reactions were filtered through a 3,000 MWCO centrifugal filter (Amicon) to remove protein and immediately used for HPLC analysis.

### ThsA NADase assay

NADase assays were performed with a 1:20 dilution of crude ThsB product or 100 nM purified 1’’-3’ gcADPR diluted into final reactions (25 mM Tris pH 7.4, 100 mM NaCl, 10% glycerol, 2 mM *β*-mercaptoethanol, 100 *μ*M NAD+ and 1 μM enzyme) and were started with the addition of enzyme. All reactions were incubated for 2 hours at 37°C. To isolate the degradation products for HPLC analysis, nucleotide synthesis reaction conditions were scaled up to 200 *μ*L reactions. Reactions were incubated with gentle shaking for 2 hr at 37°C. Following incubation, reactions were filtered through a 3,000 MWCO centrifugal filter (Amicon) to remove protein and immediately used for HPLC analysis.

### NAD+ colorimetric assay

Detection of NAD from cell lysates was performed using an NAD/NADH colorimetric assay kit (Abcam, ab65348). To generate lysates for analysis, an overnight culture of *S. aureus* RN4220 with partial or full Thoeris was diluted 1:100 and outgrown to mid-log phase (∼90 min) in BHI broth supplemented with 5 mM CaCl_2_. The culture was diluted to OD_600_ of 0.3. The culture was either infected by adding phage at MOI 1, or a prophage was induced with the addition of 1 *μ*g/ml MMC. The infected or induced cultures were grown at 37°C with shaking for 1-2 hrs. Pelleted cells were resuspended in 1X PBS with lysostaphin. After incubating cells at 37°C for 45 min, the resulting lysate was used for analysis and processed according to the manufacturers protocol.

### Nucleotide HPLC Analysis

Reaction products were analyzed using the 1460 HPLC system (Agilent) with a diode array detector at 260 nm. Sample (10 *μ*l) was loaded onto a C18 column (100 x 2.0 mm, S-3 *μ*m, 12 nm; YMC) equilibrated in 60 mM KH_2_PO_4,_ 40 mM K_2_HPO_4_ buffer. Separation was performed at a flow rate of 1.2 mL min^-1^ using a gradient program for mobile phase (acetonitrile): 0-10 min.

### Structural prediction and analysis

The amino acid sequences of Thoeris proteins or Φ80α-vir, ΦNM1, ΦNM4, ΦJ1, ΦJ2, ΦJ4, and Φ12 Major Heads were used to seed a position-specific iterative BLAST (PSI-BLAST) search of the NCBI non-redundant protein and conserved domain databases (composition-based adjustment, E-value threshold 0.01). A structure for all major head proteins was predicted using AlphaFold3 as a monomer or hexamer. Following structure determination, pairwise structural comparison of the rank 0 models was performed using PyMol. All predicted structures were compared to the solved structure of the Φ80α prohead (PDB: 6B0X). The ConSurf database was used to visualize and pinpoint conserved structural and functional features of the major heads.

### Time-lapse fluorescence microscopy

*S. aureus* cells harboring incomplete (*thsB1/B2*) or full (*thsA/B1/B2*) Sau-Thoeris were loaded onto microfluidic chambers using the CellASIC ONIX2 microfluidic system. After cells became trapped in the chamber, they were supplied with BHI medium with 5 mM CaCl_2_ under a constant flow of 5 μl h^-1^. After 1 hr, GFP-tagged Φ80α-vir was flowed through the chambers for 1 hr, before switching back to growth medium. Phase contrast images were captured at 1,000x magnification every 2 min using a Nikon Ti2e inverted microscope equipped with a Hamamatsu Orca-Fusion SCMOS camera and the temperature-controlled enclosure set to 37°C. GFP was imaged using a GFP filter set using an Excelitas Xylis LED Illuminator set to 2% power, with an exposure time of 100 ms. Images were aligned and processed using the NIS Elements software.

### Generation and isolation of escaper bacteriophages

Overnight cultures of *S. aureus* RN4220 were diluted 1:100 and outgrown at 37°C with shaking for 1 hr, infected with Φ80α-vir (MOI 1) for 20 min. Cultures were allowed to lyse for 3 hr before pelleting debris and sterile-filtering the supernatant to obtain phage. 100 μL of RN4220 overnight cultures harboring Sau-Thoeris were infected with a high titer mutant phage library in BHI soft agar and then plated. All plaques were collected and the soft-agar infection was repeated five times. After the fifth passage at 37°C overnight, individual phage plaques were picked from the top agar and resuspended in 50 μL of BHI liquid medium. Phage lysates were further purified over two rounds of passaging on RN4220 harboring Sau-Thoeris. Genomic DNA from high titer phage stocks was extracted using previously described methods (Jakociune and Moodley, 2018) and was submitted to SeqCenter for whole genome sequencing and assembly.

### Cobalt enrichment of ThsB complex

His-tagged ThsB1 or ThsB2 were each expressed with the complementary 3xFLAG-tagged ThsB. After expression of the ThsB proteins with 1 mM IPTG, cells were either infected with Φ80α-vir (MOI 10) for 20 min or ΦNM1 prophage was induced by the addition of 1 μg/mL mitomycin C for 1 hr 30 min.

Where indicated, the ThsB proteins were co-expressed with Mhp from a plasmid. The resulting cells were collected by centrifugation and resuspended in 25 mM Tris pH 7.4, 100 mM NaCl, 10% glycerol, 2 mM *β*-mercaptoethanol, 5 mM MgSO_4_ with lysostaphin, DNase1 and EDTA-free protease inhibitor cocktail. The cells were lysed at 37°C with shaking for 30 min before brief sonication. Lysates were clarified by ultracentrifugation and applied to cobalt affinity resin (∼0.2 mg). After binding, the resin was washed six-ten times with lysis buffer prior to elution with lysis buffer containing 200 mM imidazole. Eluted proteins were visualized by SDS-PAGE and western blot using anti-His6 and anti-3xFLAG antibodies.

### Phylogenetic analysis of Thoeris from *S. aureus*

Bioinformatically predicted Thoeris systems in S. aureus were identified in Doron et al., 2018 (Doron *et al*., 2018). Unique Thoeris systems were identified by analyzing the protein sequences of these predicted systems in Geneious Prime 2024.0.5.

### Statistical analysis

All statistical analyses were performed using GraphPad Prism v9.5.1. Error bars and number of replicates for each experiment are defined in the figure legends. Comparisons between groups for viral titer, gene expression, colony-forming units, and NAD+ concentration were analyzed by unpaired parametric t-test, two-tailed with no corrections.

## Acknowledgements

We would like to thank the members of the Marraffini laboratory for constructive feedback and encouragement. LAM is an investigator of the Howard Hughes Medical Institute. CBF is supported by the National Science Foundation Graduate Research Fellowship under Grant 1946429. DVB is supported by an NIH Ruth L. Kirschstein NRSA F30 Individual Predoctoral Fellowship (F30AI157535) and an NIH Medical Scientist Training Program grant (T32GM152349) to the Weill Cornell/Rockefeller/Sloan Kettering Tri-Institutional MD-PhD Program. We would also like to thank the Rockefeller University Proteomics core for their assistance with mass spectrometry.

## Author contributions

Project was conceived by CGR. Experiments were designed by CGR, CBF, DVB, and LAM. DVB performed cloning and initial testing of the Sau-Thoeris system. CGR and CBF conducted and analyzed all other experiments. The paper was written by CGR, CBF, and LAM, with input from DVB.

## Data availability

Source data are provided with this paper. Any additional data from this study are available from the lead contact upon request.

## SUPPLEMENTARY FIGURE LEGENDS

**Figure S1.**
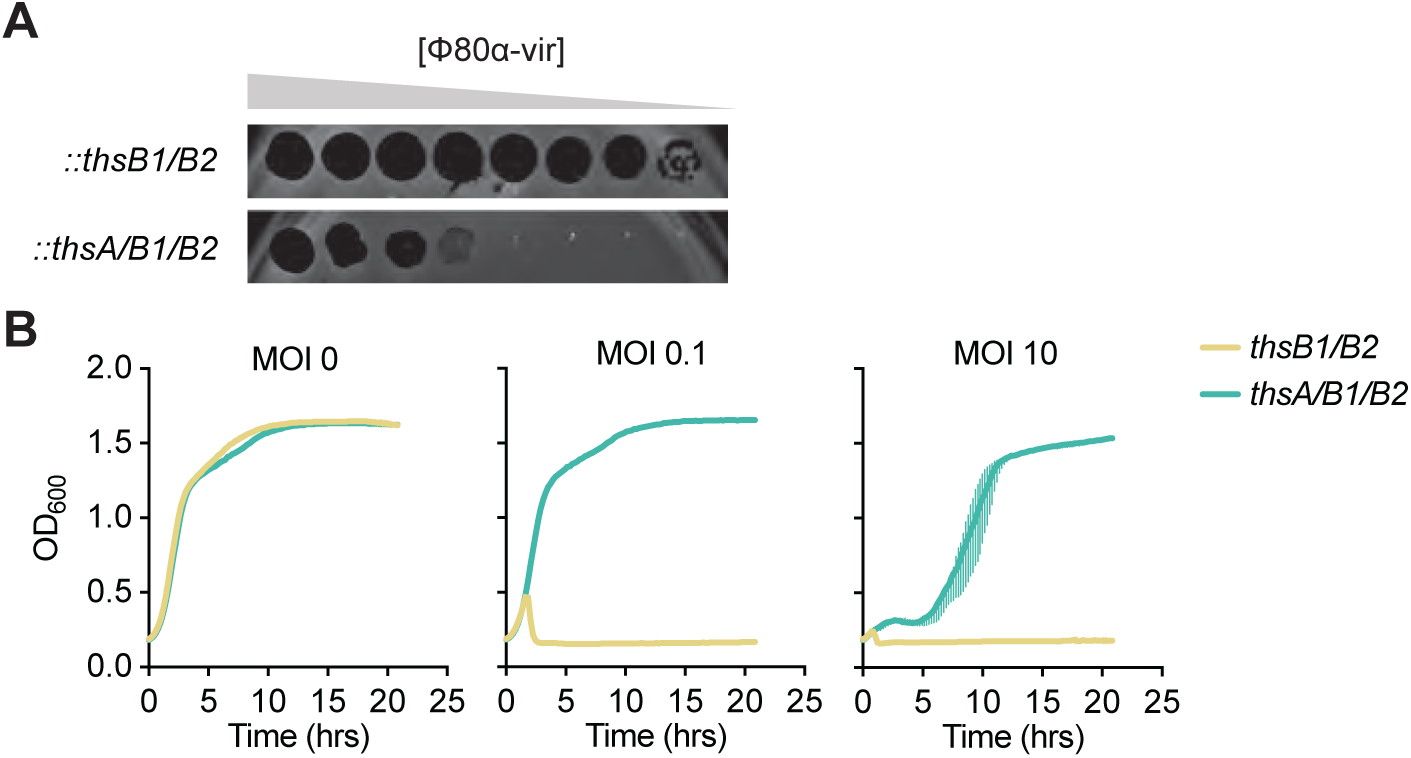
Characterization of Sau-Thoeris defense. **(A)** Tenfold serial dilutions of Φ80α-vir on lawns of *S. aureus* RN4220 carrying either an incomplete (*thsB1/B2*) or full (*thsA/B1/B2*) Thoeris operon in a chromosomal location. **(B)** Growth of *S. aureus* RN4220 harboring plasmids carrying either an incomplete (*thsB1/B2*) or full (*thsA/B1/B2*) Thoeris operon, determined as the OD_600_ of the cultures after infection with Φ80α-vir at MOI 0, 0.1 or 10. Mean of +/- S.D. of three biological replicates is reported.

**Figure S2.**
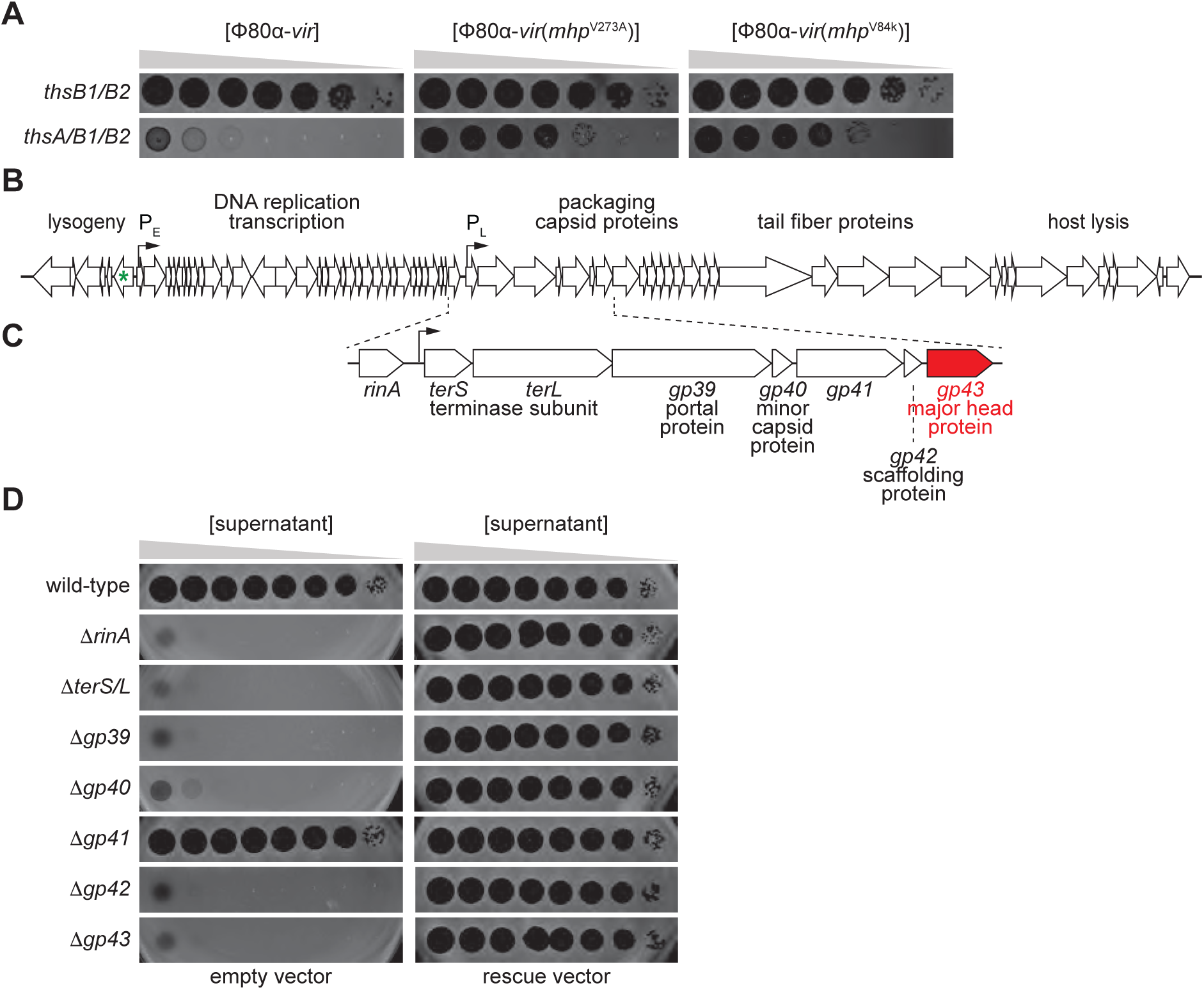
Major head protein is required for Sau-Thoeris activation. **(A)** Schematic of ΦNM1 genome, with expansion of the operon for packaging the genome and assembly of phage particles. P_E_ and P_L_ are promoters responsible for the expression of early and late viral genes, respectively. The green asterisk indicates the nonsense mutation prevents ΦNM1 lysogeny, converting it into ΦNM1γ6, a purely lytic phage. Regions of the genome involved in different stages of the viral lytic cycle are indicated. **(B)** Genes of the ΦNM1 genome required for capsid biogenesis. **(C)** Tenfold serial dilutions of Φ80a-vir phage carrying wild-type or V273A *mhp* alleles on lawns of *S. aureus* RN4220 harboring plasmids carrying either an incomplete (*thsB1/B2*) or full (*thsA/B1/B2*) Thoeris operon. **(D)** Tenfold serial dilutions of lysates obtained after MMC induction of different ΦNM1 lysogens carrying deletions on genes involved in packaging the genome and assembly of phage particles on lawns of *S. aureus* RN4220 harboring an empty vector control, or plasmids expressing the genes deleted in the prophages.

**Figure S3.**
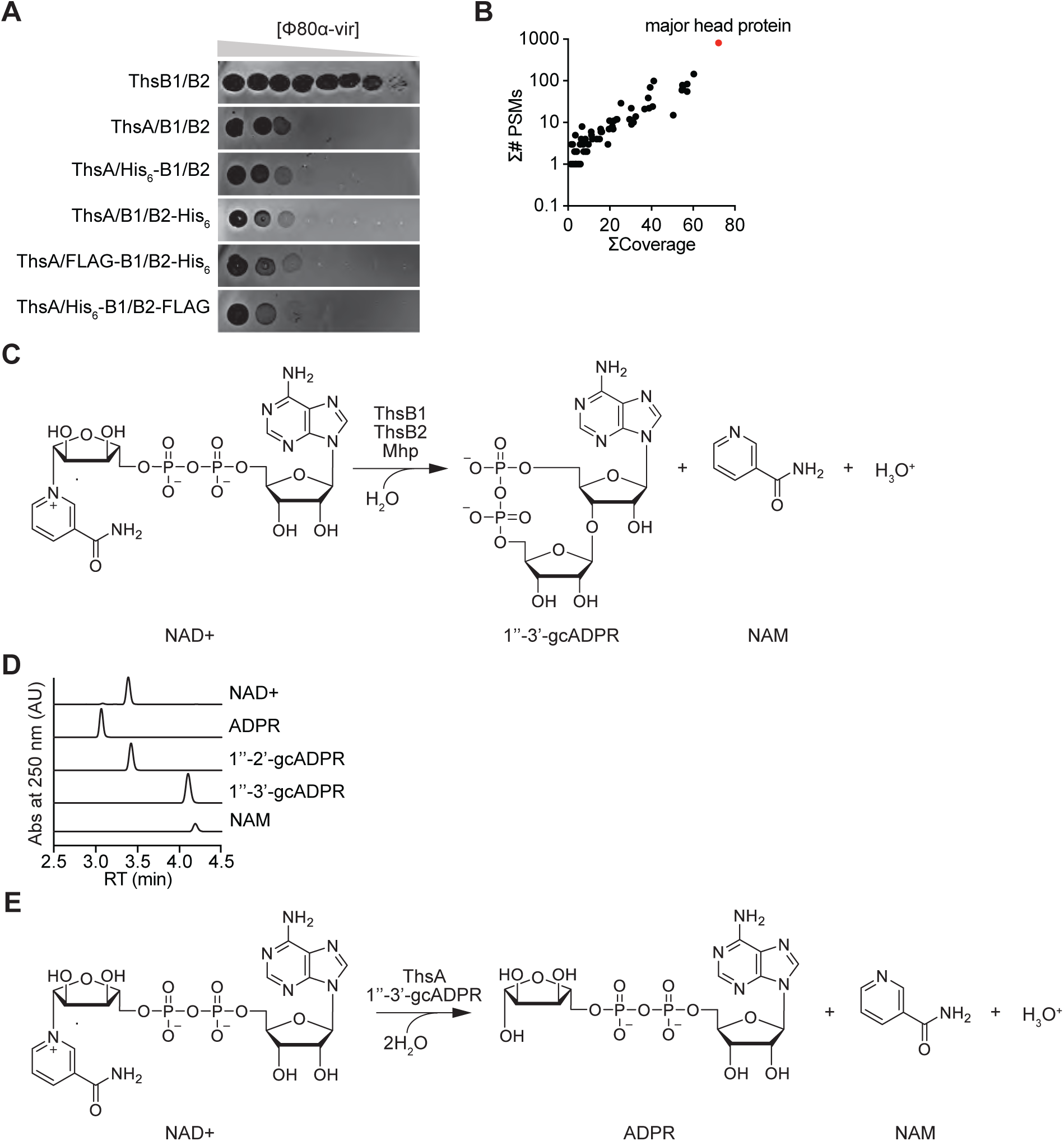
ThsB1-ThsB2-Mhp copurification assays. **(A)** Tenfold serial dilutions of Φ80a-vir phage on lawns of *S. aureus* RN4220 harboring plasmids carrying either untagged or hexahystidyl- or FLAG-tagged versions of ThsB1 or ThsB2, along with untagged ThsA. **(B)** LC-MS/MS identification of ∼35 kDa protein isolated with ThsB1 and ThsB2 from infected *S. aureus* RN4220 cells shown in Figure 3B. Gel slices (n=3) were subjected to reduction, alkylation, and in-gel digestion. Peptides were extracted before being injected for LC-MS/MS analysis. Search results against the staphylococcal phage ΦNM1 proteome are presented as the percentage of sequence coverage (ΣCoverage) vs the indication of unique identified peptides (ΣPSMs). Data for the phage Mhp is shown in red. **(C)** Cyclization reaction of oxidized nicotinamide adenine dinucleotide (NAD+) mediated by the ThsB1/B2/Mhp complex, which yields 1”-3’-glyocyclic ADP-ribose (1”-3’-gcADPR) and nicotinamide (NAM). **(D)** HPLC analysis of different commercially available chemicals used in this study. **(E)** Hydrolysis reaction of oxidized nicotinamide adenine dinucleotide (NAD+) mediated by ThsA when activated by 1”-3’-gcADPR, which yields ADP-ribose (ADPR) and nicotinamide (NAM).

**Figure S4.**
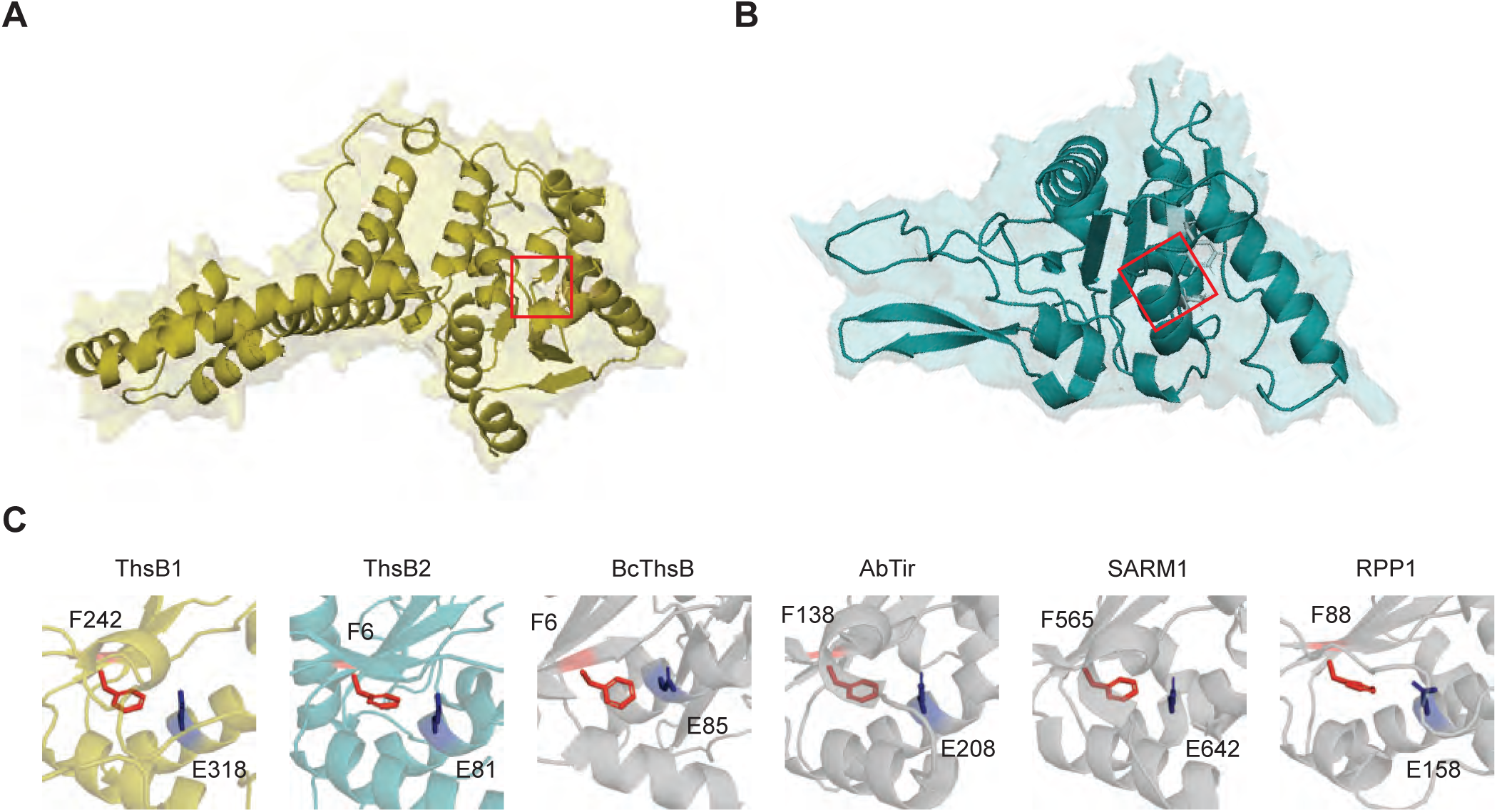
Structural analysis of ThsB1 and ThsB2. **(A)** Structure of ThsB1 generated by AlphaFold3, with the putative active site marked with a red square. **(B)** Same as **(A)** but for ThsB2. **(C)** Structural comparison of the predicted active sites of ThsB1 and ThisB2 (red squares in panels A and B, respectively) with the experimentally characterized active sites of different TIR proteins: *Bacillus cereus* ThsB (BcThsB; PDB: 6LHY), *Acinetobacter baumannii* TIR domain (AbTir; PDB: 7UXU), human sterile alpha and TIR motif containing preotein 1 (SARM1; PDB: 6O0R) and *Arabidopsis thaliana* resistance protein RPP1 (PDB: 7DFV).

**Figure S5.**
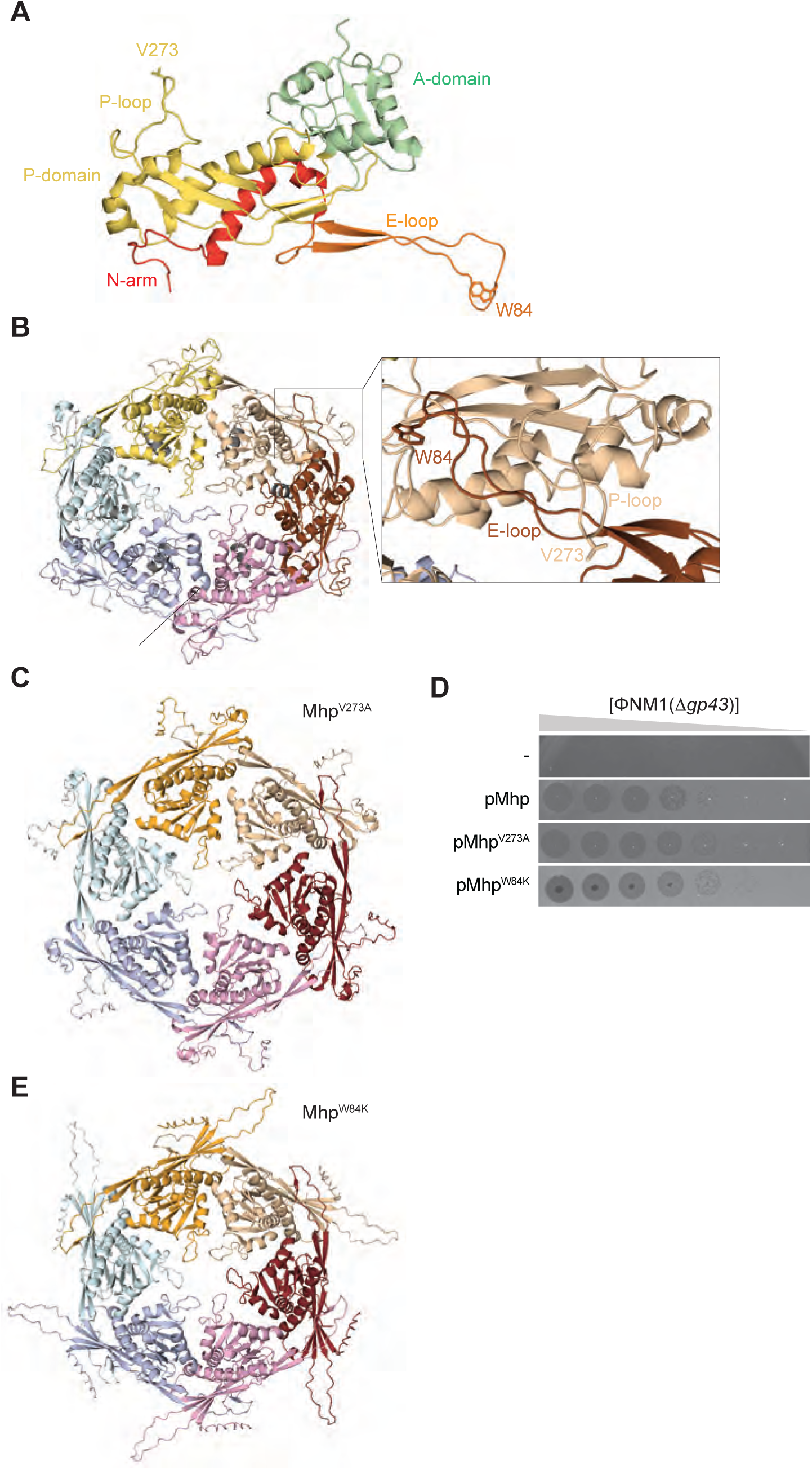
Importance of Mhp hexameric form for Sau-Thoeris activation. **(A)** Structure Φ80α Mhp monomer (PDB: 6B0X), colored according to the different structural domains (red, N-arm; yellow, P domain and loop; green, A-domain; orange, E-loop). **(B)** Structure of the hexameric form of Φ80α major head protein (PDB: 6B0X), with one capsomer interface highlighted. The position of the residues which mutations led to immune evasion, V273 within the P-loop (tan) and W84 within the E-loop (brown), are shown. **(C)** Alphafold3 prediction of the hexameric form of Φ80α Mhp^V273A^. **(D)** Tenfold serial dilutions of lysates obtained after MMC induction of a ΦNM1(1*gp43*) lysogen on lawns of *S. aureus* RN4220 harboring either an empty vector control (-), or plasmids expressing wild-type, V273A or W84K Mhp. **(E)** Sames as **(C)** but for Φ80α _MhpW84K._

**Figure S6.**
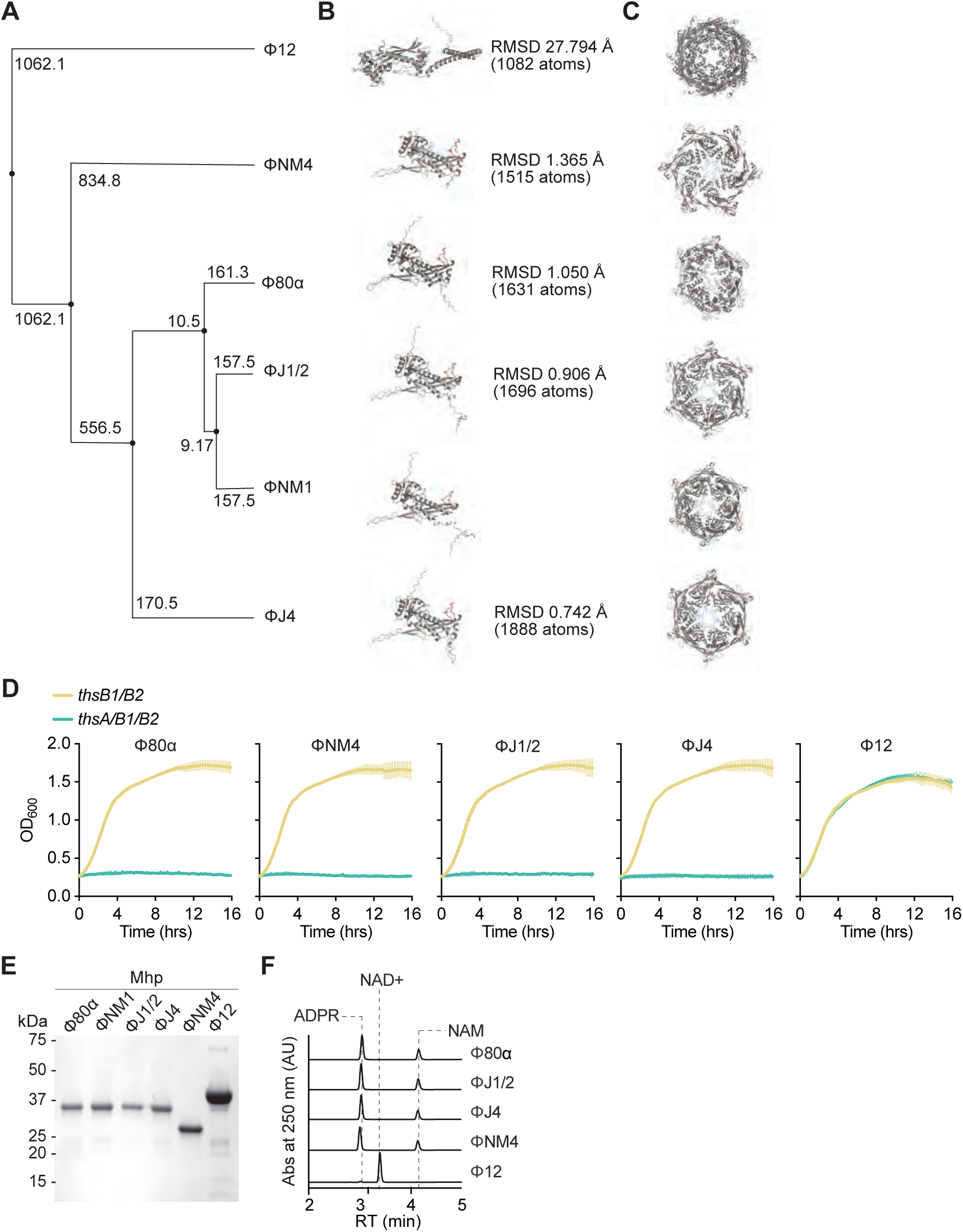
Analysis of the Sau-thoeris activating properties of Mhp from different staphylococcal phages. **(A)** Phylogenetic tree of the major head proteins used in this study, generated using EMBL-EBI Muscle multiple sequence alignment tool. The branch numbers indicate genetic distance from ΦNM1 Mhp calculated by the software. **(B)** Alphafold3 prediction of the monomeric form of Mhp encoded by the different phages used in this study. Structural distance is shown as the RMSD of the different Mhps compared to ΦNM1 Mhp. **(C)** Alphafold3 prediction of the hexameric form of Mhp encoded by the different phages used in this study. **(D)** Growth of *S. aureus* RN4220 harboring plasmids carrying either an incomplete (*thsB1/B2*) or full (*thsA/B1/B2*) Thoeris operon in the presence of a second plasmid expressing Mhp from different staphylococcal phages, determined as the OD_600_ of the cultures after addition of IPTG. The values at the end of the curves, at 16 hours, were used to meke the bar graphs shown in Figure 6A. Mean of +/- S.D. of three biological replicates is reported. **(E)** Coomassie Blue-stained SDS-PAGE of Mhp proteins, purified from staphylococci harboring pMhp plasmids using PEG-enrichment. Protein molecular weight (kDa) markers are shown. **(F)** HPLC analysis of the products resulting from the incubation of purified His_6_-ThsA with the products of the reactions shown in Figure 6C, obtained after mixing ThsB2-His_6_, ThsB2^F6A^-His_6_, NAD+ purified Mhp from different staphylococcal phages. Retention times (RT) of reactants and products are marked by dotted lines.

## SUPPLEMENTARY METHODS TABLES

**Supplementary Methods Table 1.**
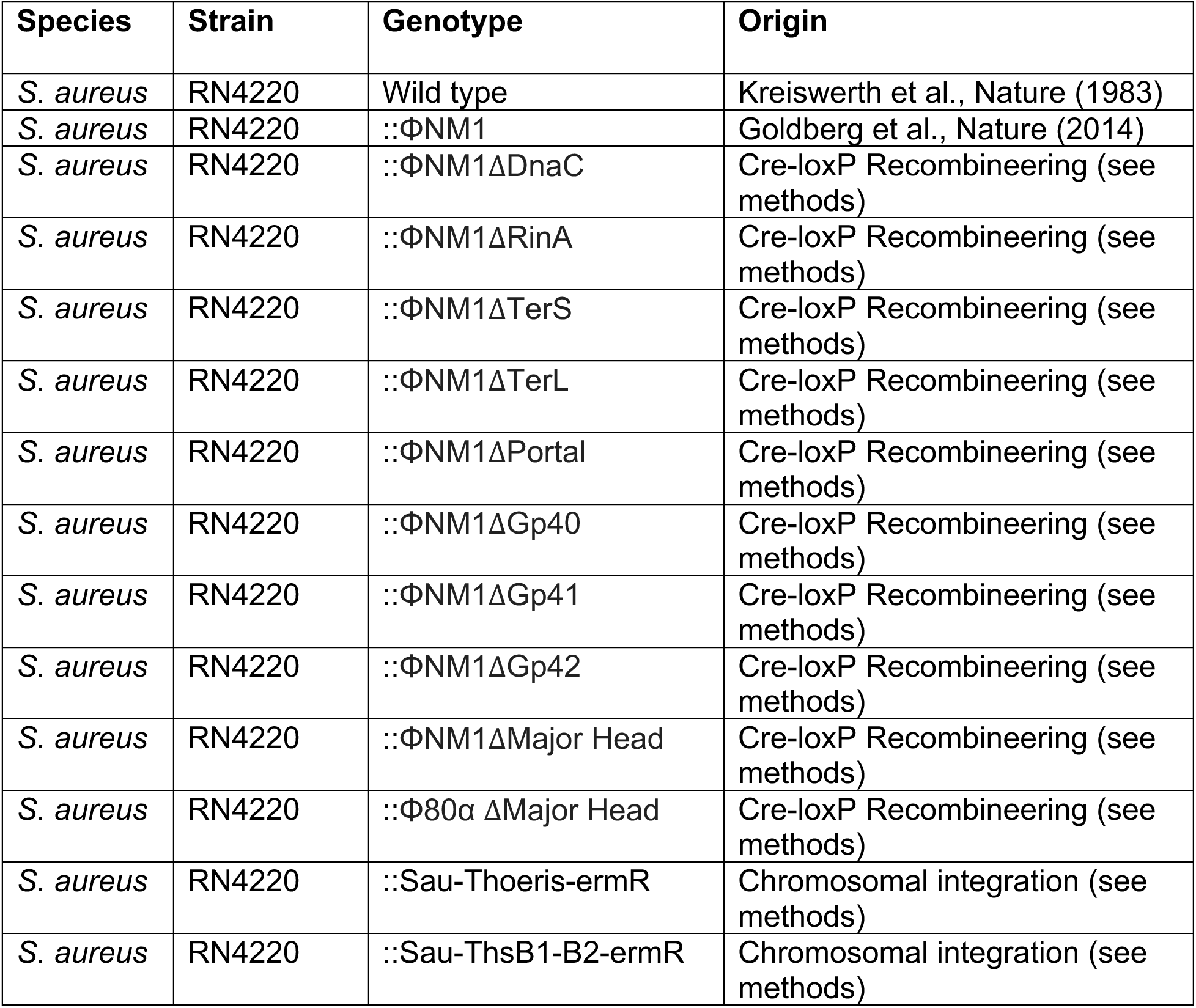
Bacterial strains used in this study.

**Supplementary Methods Table 2.**
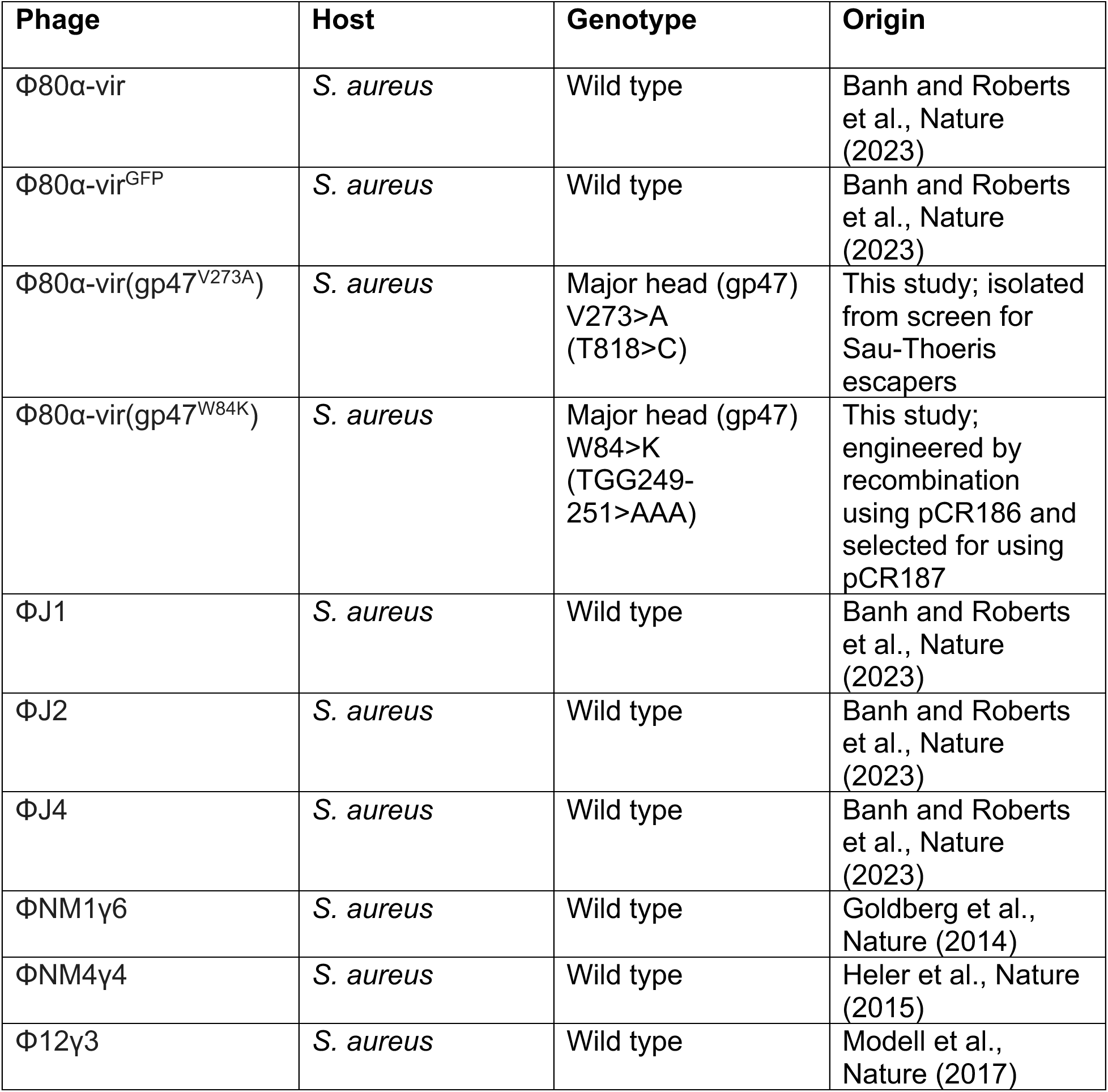
Phages used in this study.

**Supplementary Methods Table 3.**
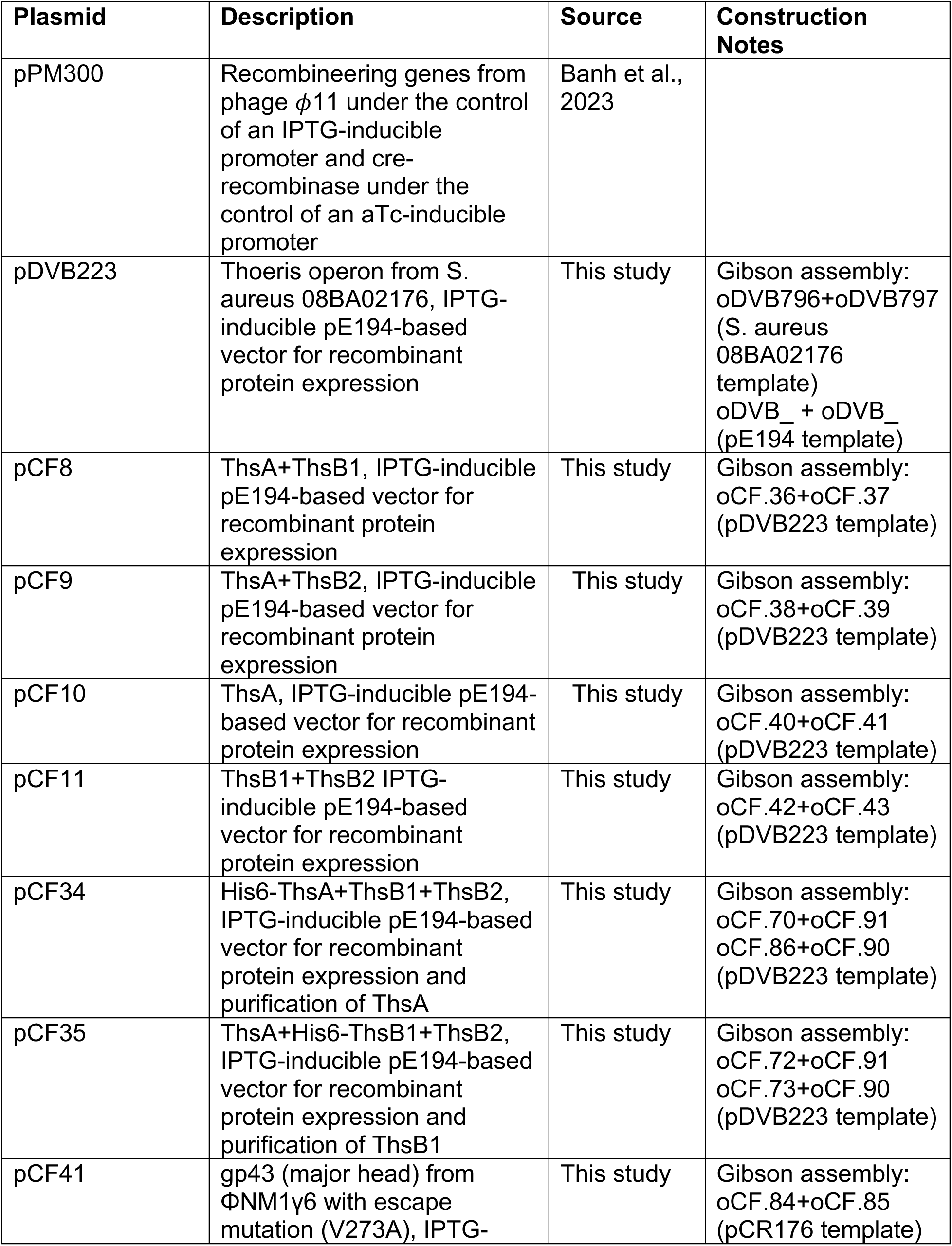
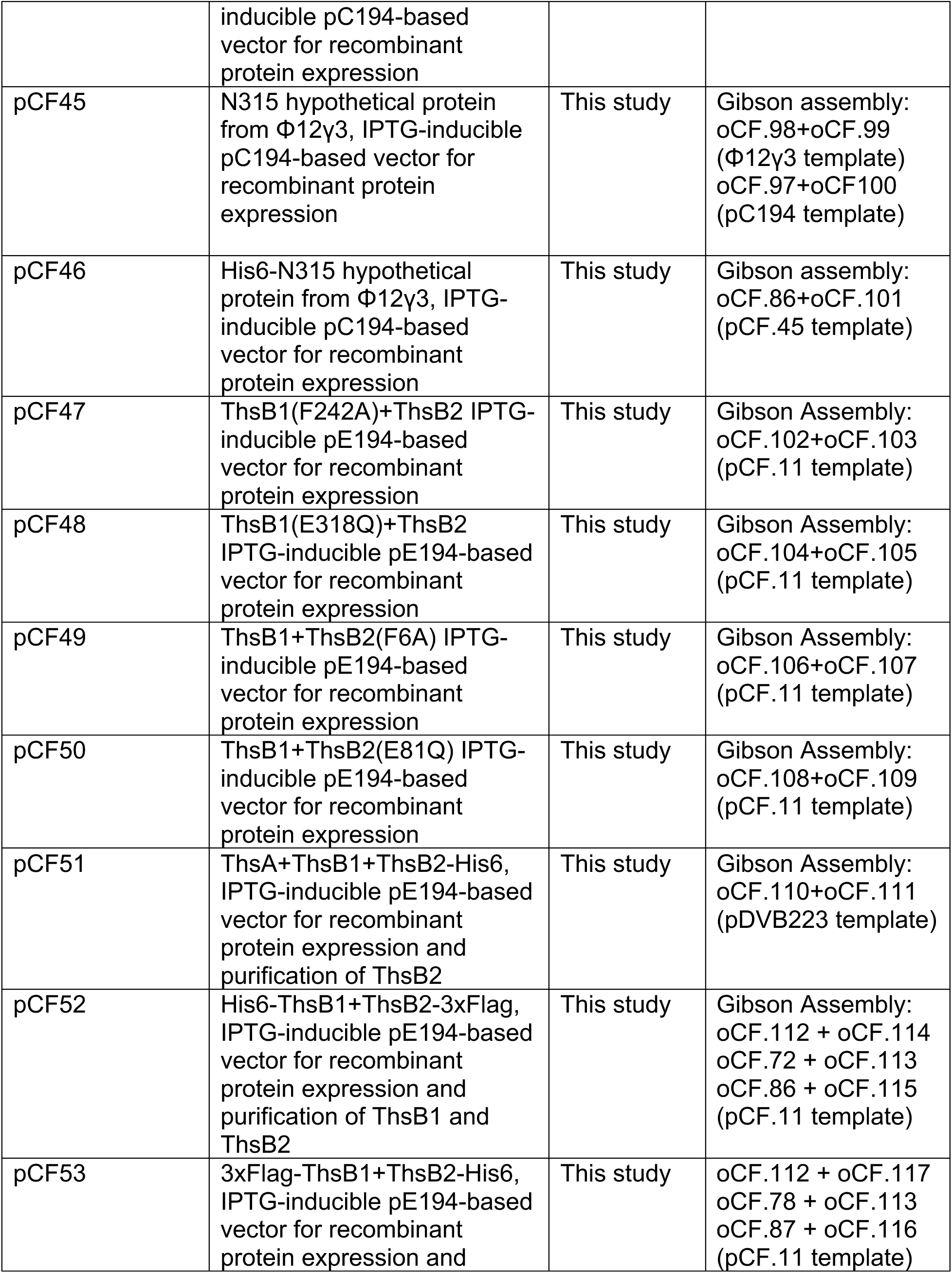
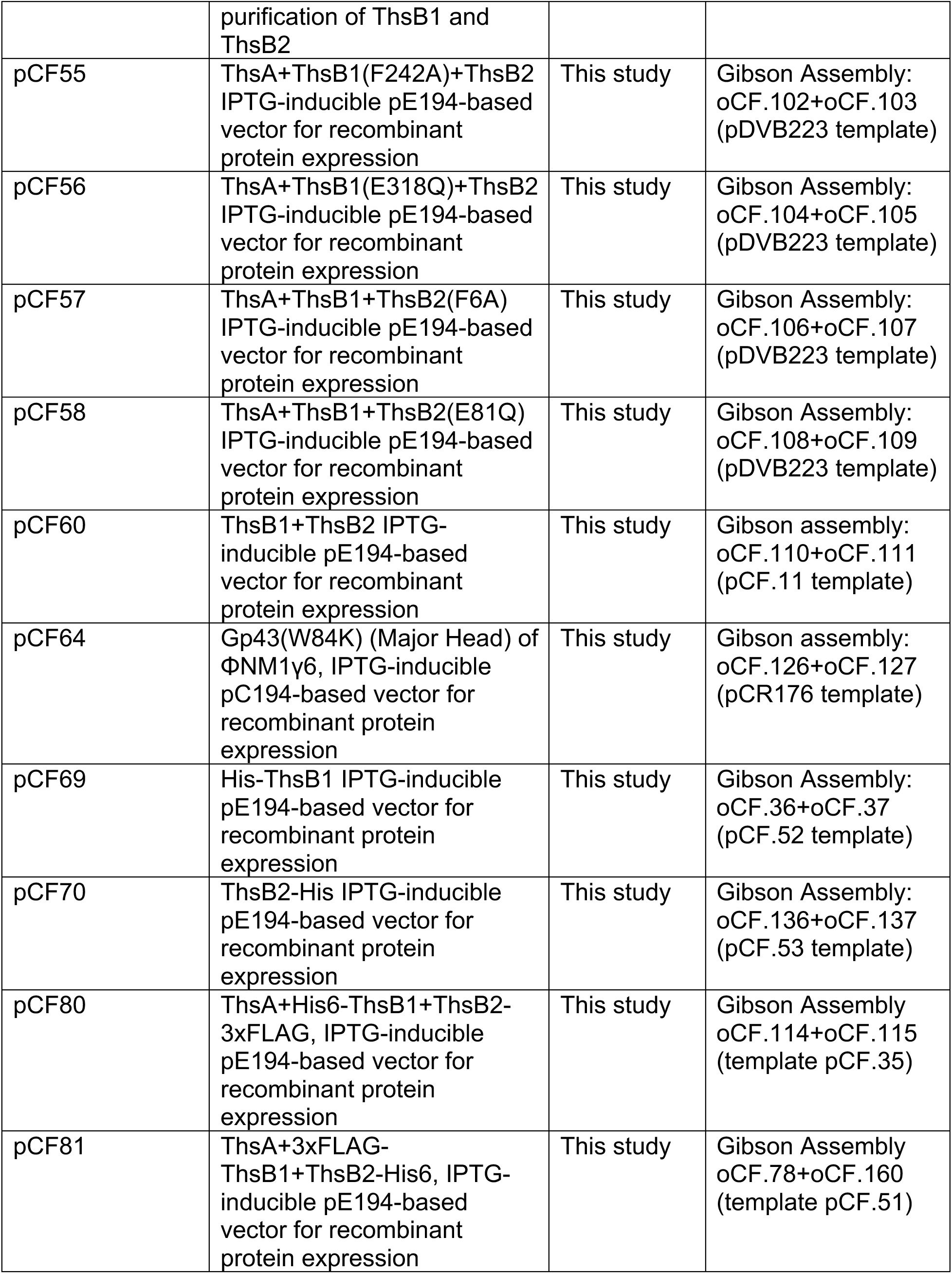
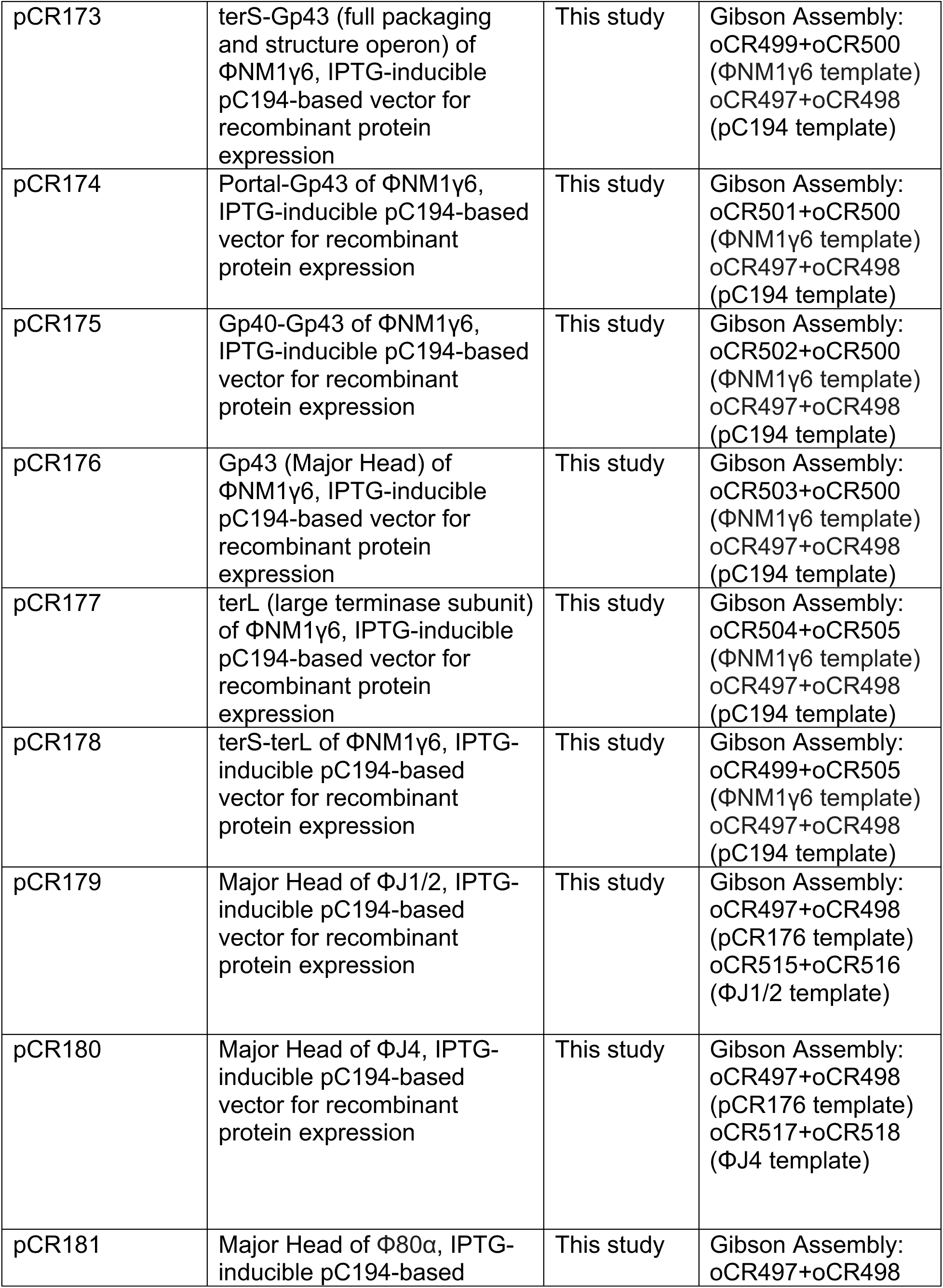
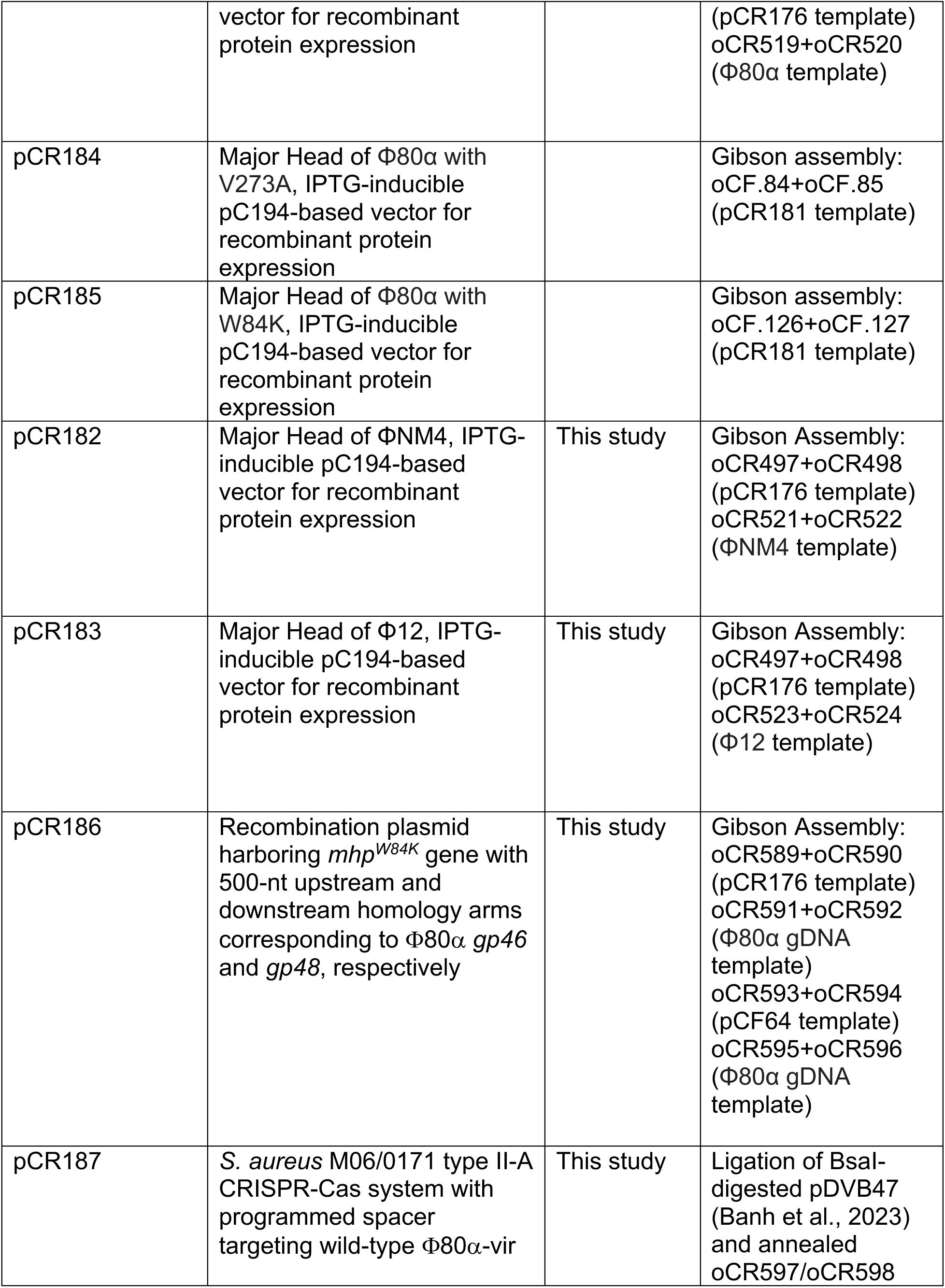
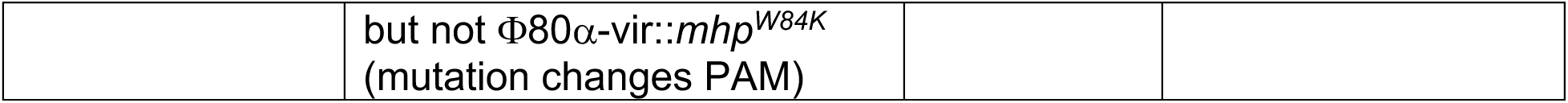
Plasmids used in this study.

**Supplementary Methods Table 4.**
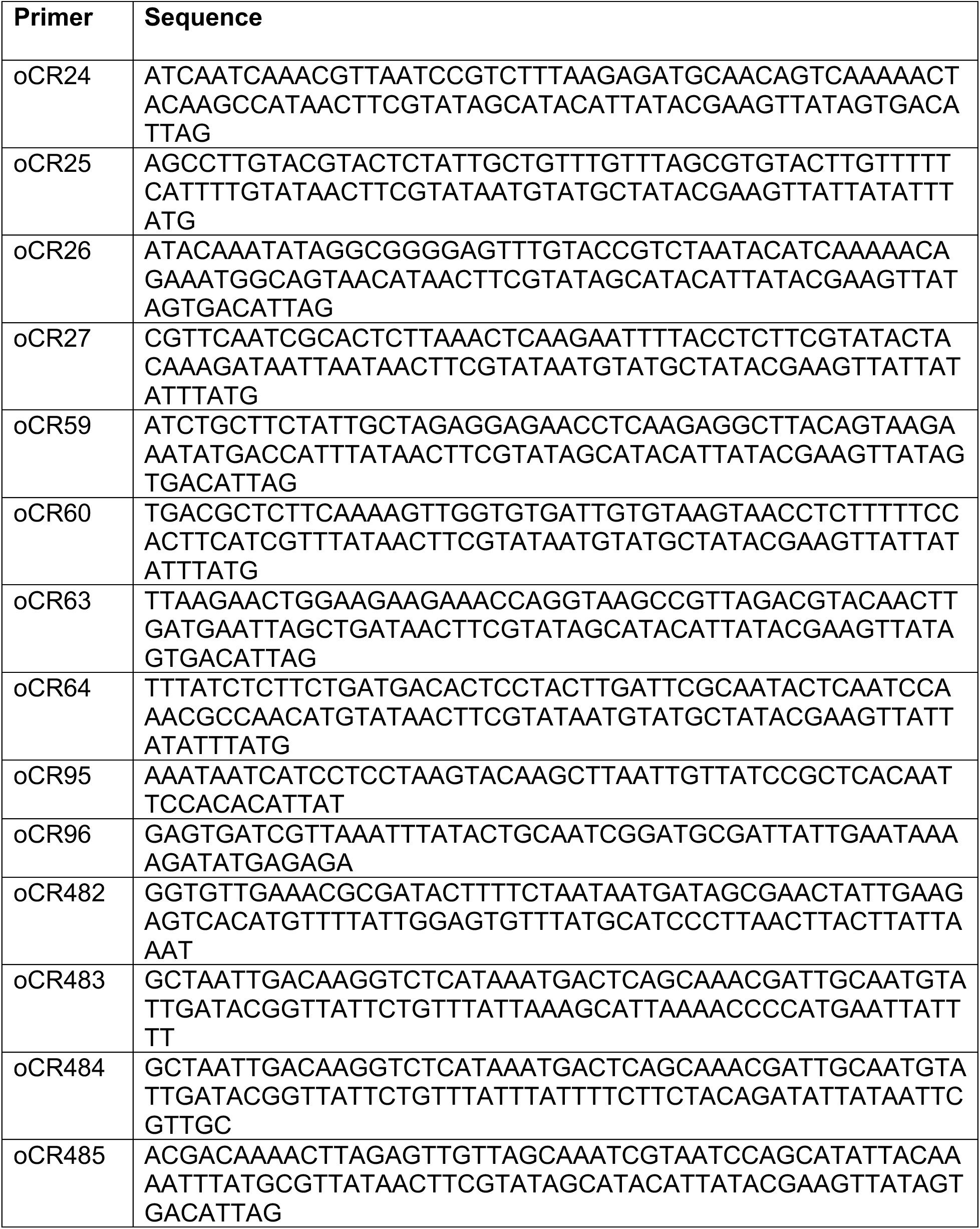
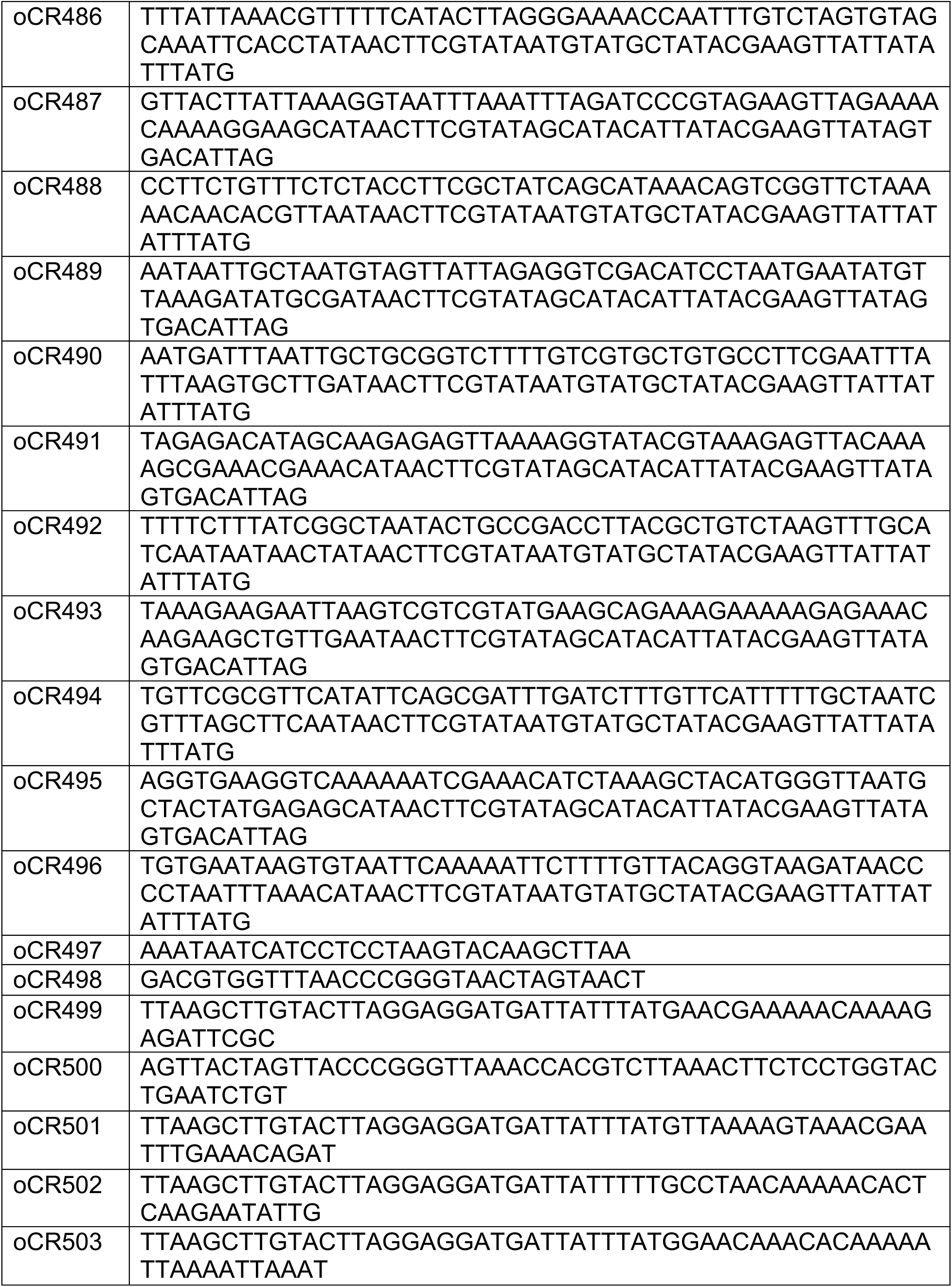
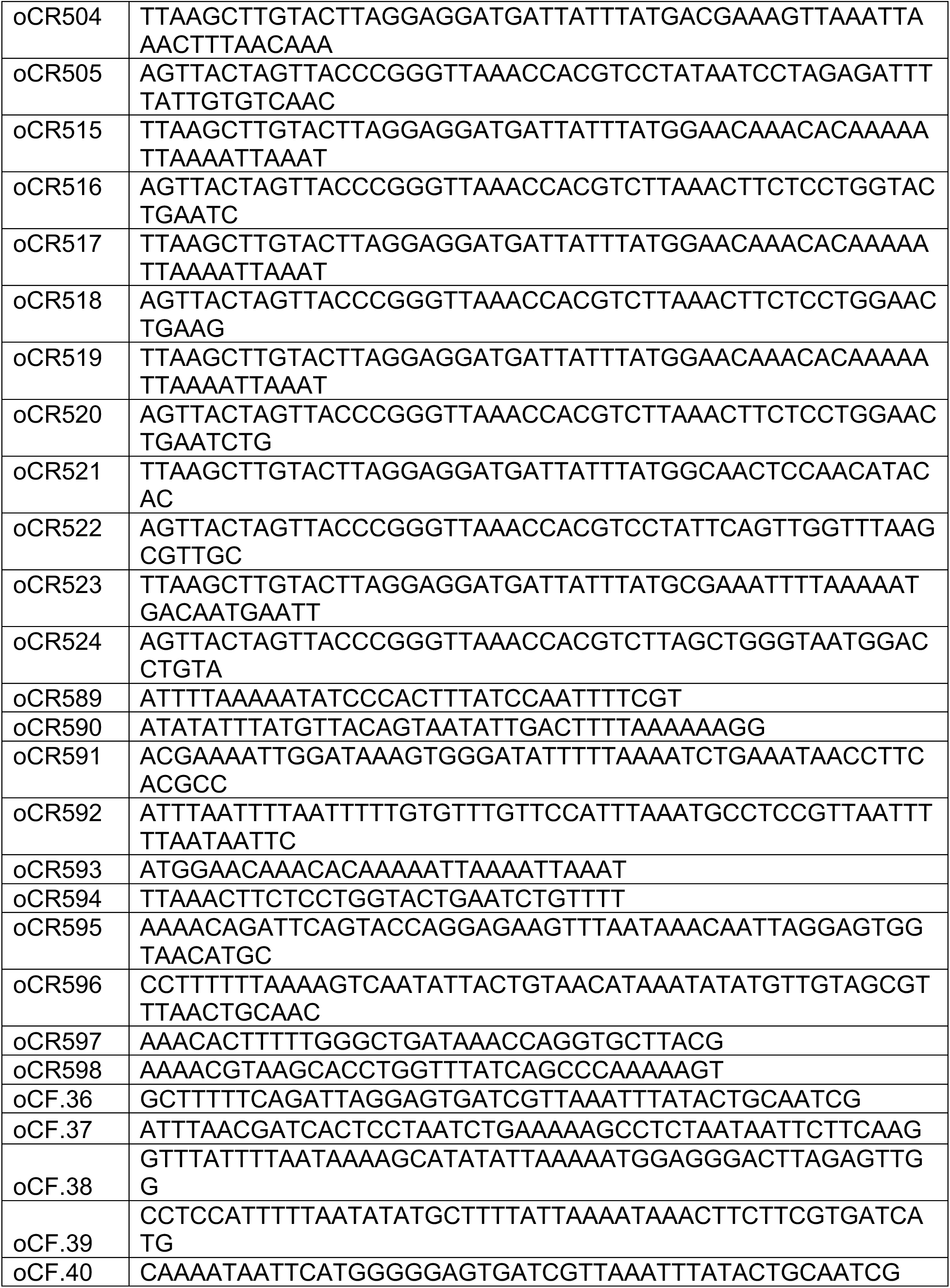
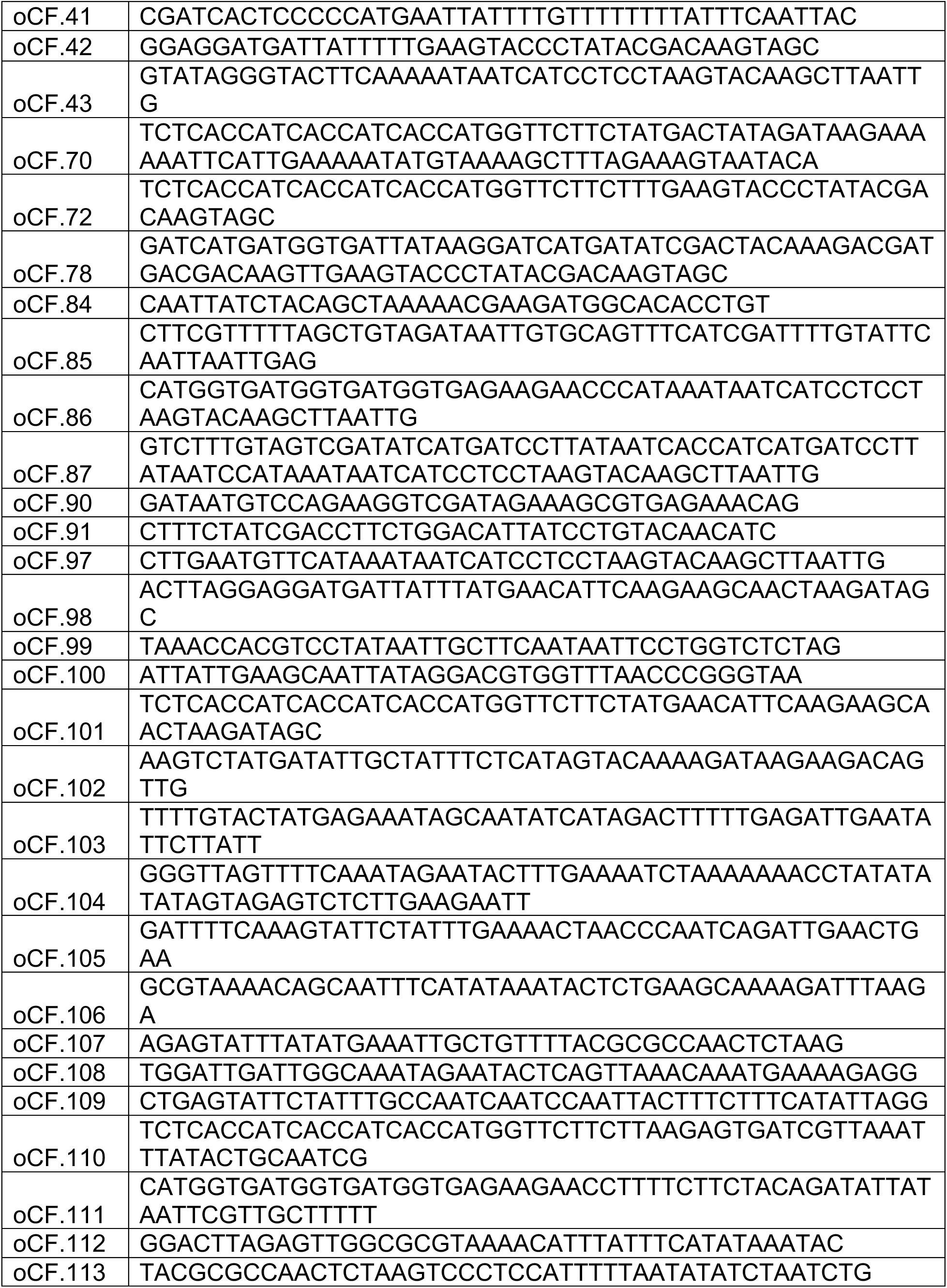
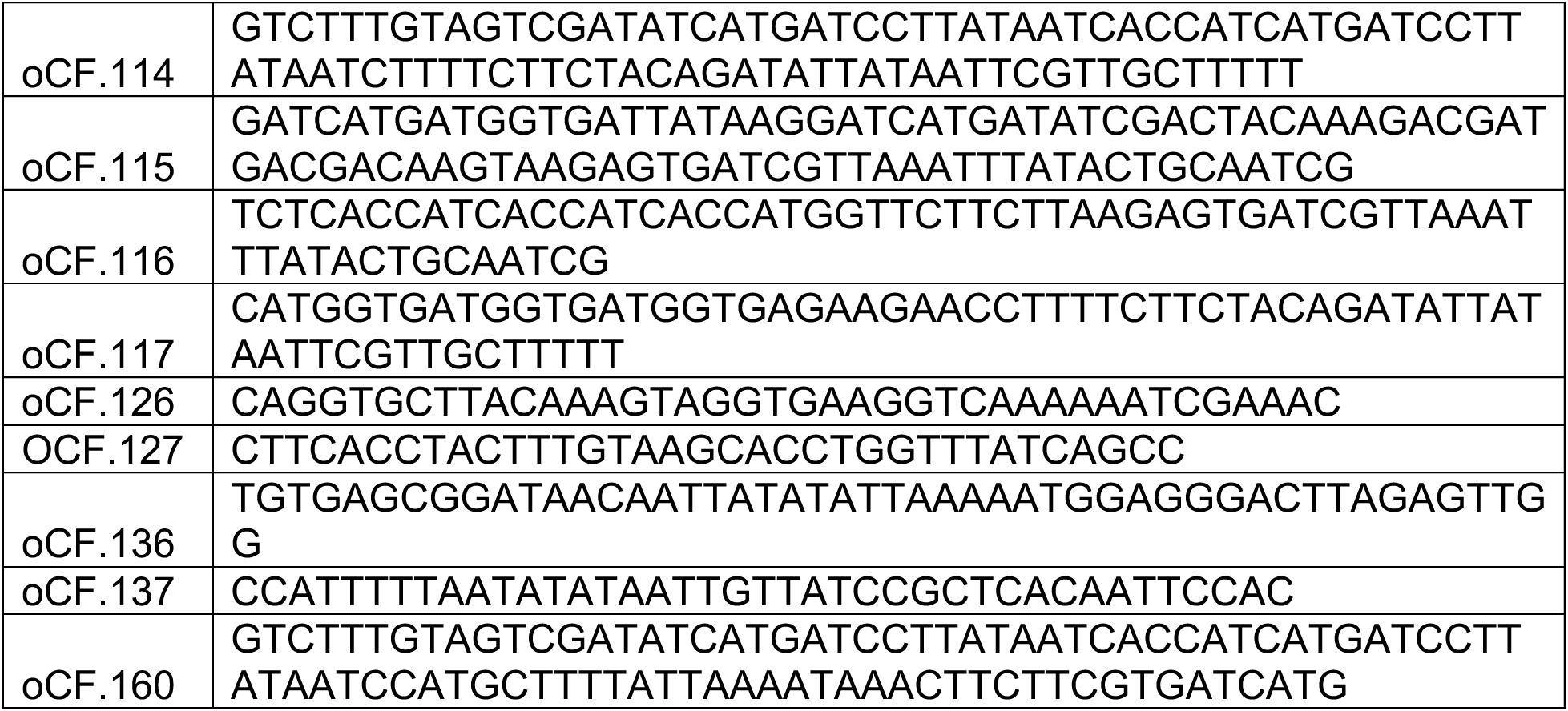
Oligonucleotide primers used in this study.

**Supplementary Sequences 1.**
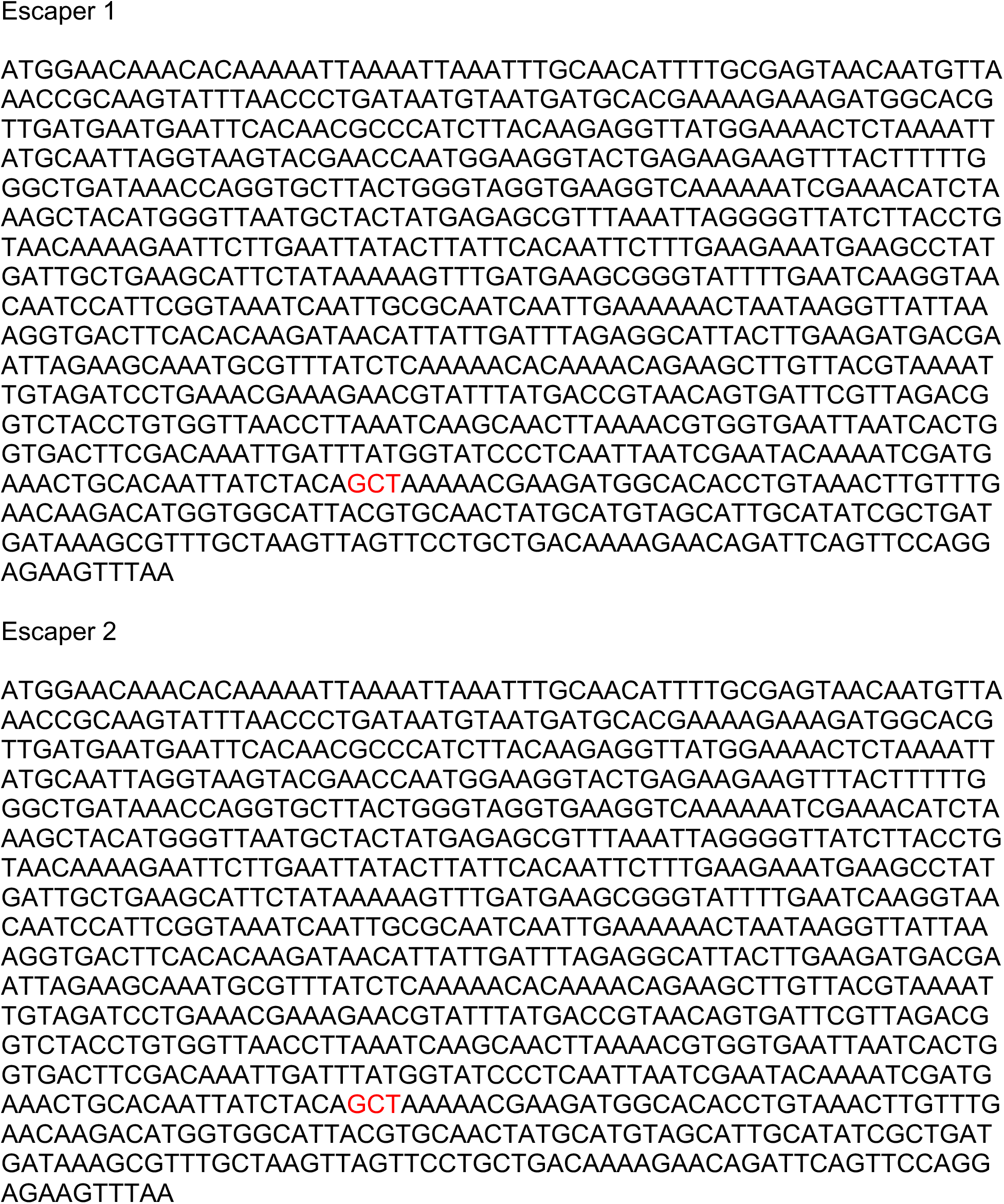
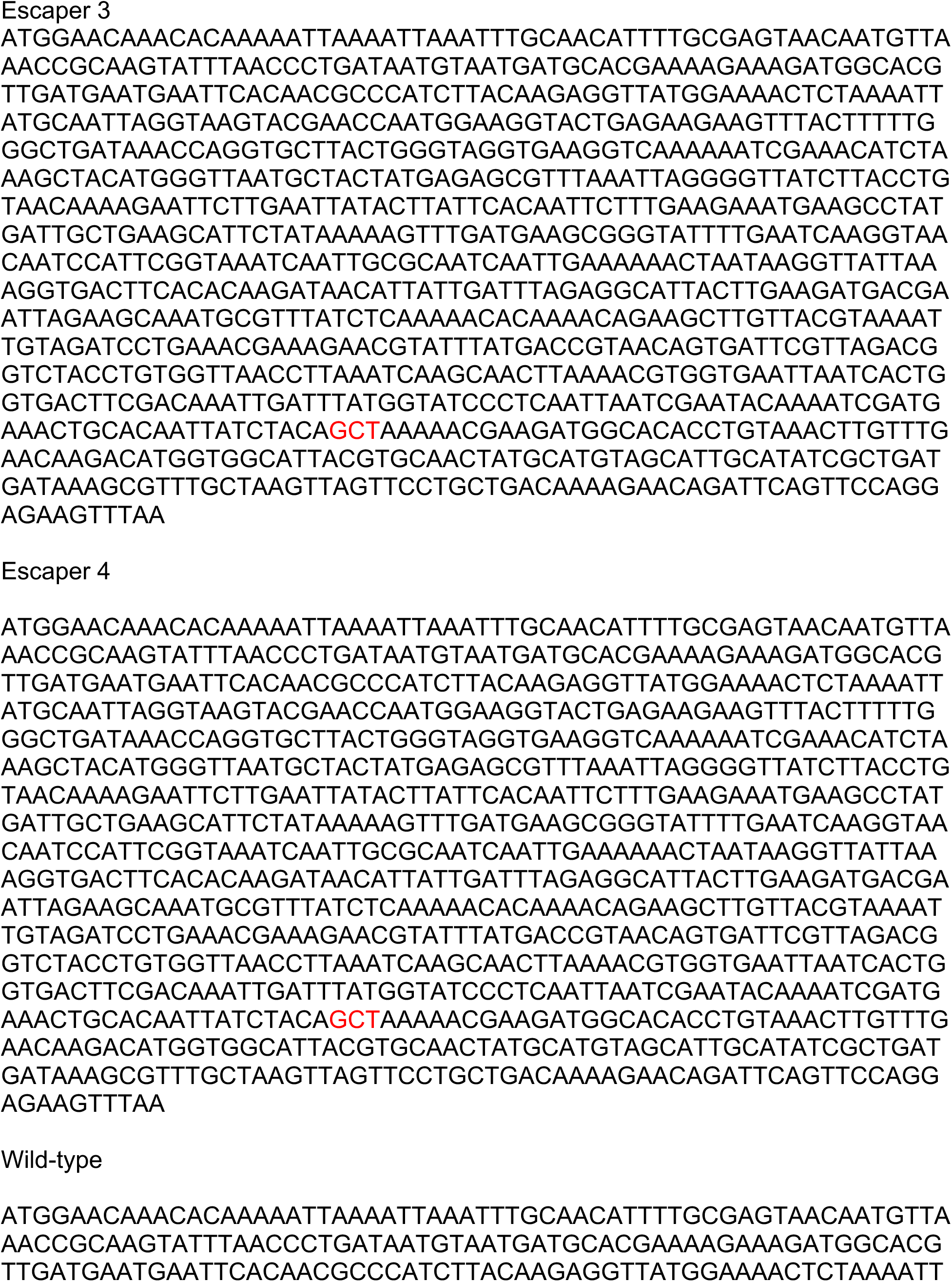
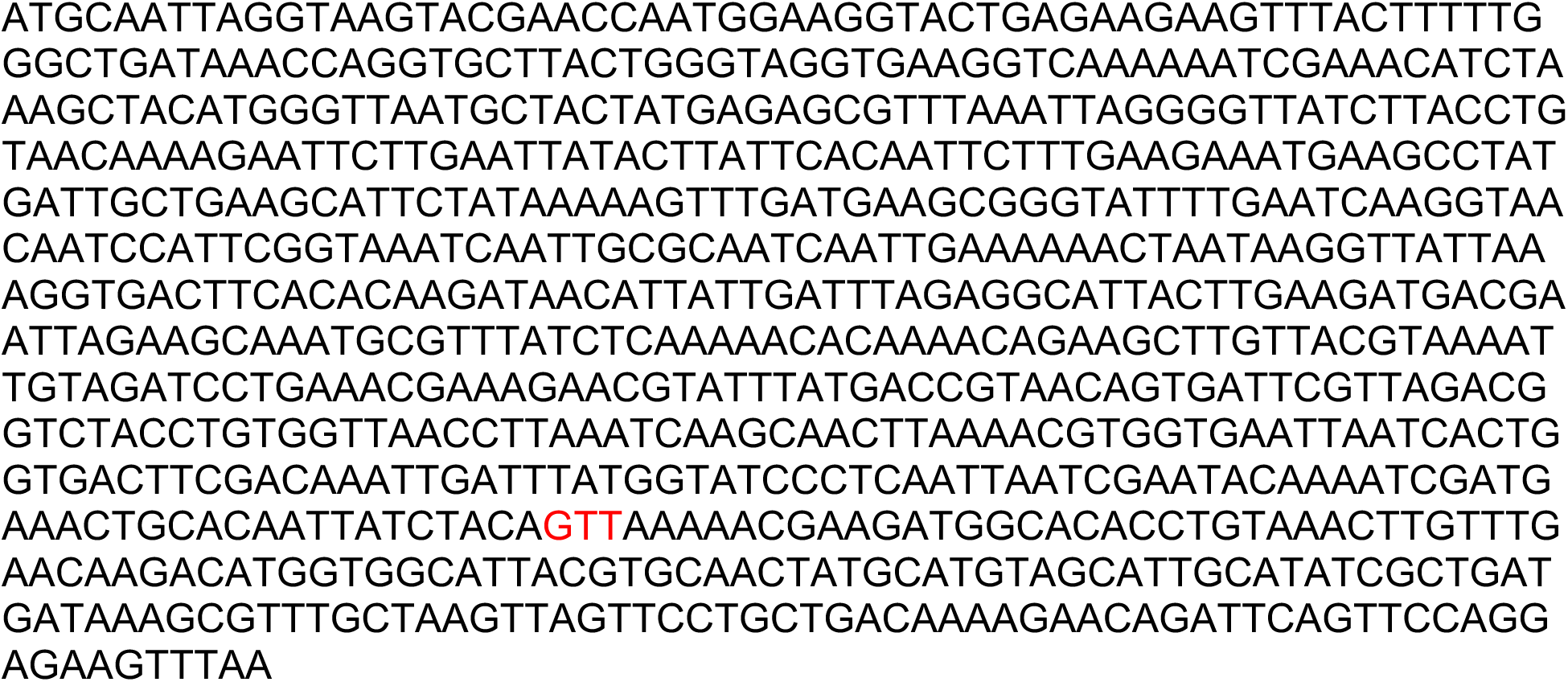
DNA sequences of *mhp* genes from escaper Φ80α-vir phages that avoid Sau-Thoeris immunity. The sequence of the codon for residue V273 is shown in red.

